# Human coronavirus nucleocapsid proteins have disparate innate immune evasion abilities

**DOI:** 10.64898/2026.07.14.738543

**Authors:** Noga Sharlin, Rory P. Mulloy, Madeline Day, Jennifer A. Corcoran

## Abstract

During infection, coronaviruses produce abundant double-stranded RNA (dsRNA) which can induce antiviral innate immune responses such as the interferon, 2’-5’-oligoadenylate synthetase (OAS)/RNase L, and protein kinase R (PKR) pathways. Coronaviruses must antagonize these dsRNA responses for successful replication. The SARS-CoV-2 nucleocapsid (N) protein plays a central role in evasion of dsRNA responses, interacting with dsRNA to block interferon-β production and the activation of OAS/RNase L and PKR. Despite intensive study of SARS-CoV-2 N, our understanding of the innate immune evasion abilities of N proteins produced by other human coronaviruses (HCoVs) remains incomplete. Here, we provide a comprehensive comparison of HCoV N proteins expressed in a human lung cell line and show that their abilities to block dsRNA-induced innate immune responses differ. Highly pathogenic HCoV N proteins inhibited the production of interferon-β mRNA and activation of OAS/RNase L, while common cold HCoV N proteins did not. While most HCoV N proteins inhibited PKR phosphorylation, HCoV-OC43 N did not, an observation that correlated with high levels of PKR activation observed during HCoV-OC43 infection. The ability of HCoV N proteins to antagonize PKR required colocalization with dsRNA, yet the overall decrease in PKR phosphorylation mediated by N was not due to sequestration of dsRNA away from PKR. Rather, N colocalized with dsRNA and PKR at dsRNA-induced foci (dRIFs) and inhibited PKR phosphorylation within dRIFs. Sarbecovirus N proteins also relocalized PKR to G3BP1 foci, suggesting that these N proteins can inhibit PKR using two distinct mechanisms. Collectively, our work reveals an unexpected level of functional and mechanistic diversity among the innate immune evasion abilities of human coronavirus N proteins. These findings challenge existing presumptions that observations made in one coronavirus can be extrapolated to others, because even conserved essential proteins such as N can exhibit considerable functional heterogeneity.

## INTRODUCTION

Cytosolic double-stranded RNA (dsRNA) is a hallmark of viral infection. Diverse viruses, including dsRNA, DNA, and positive-sense single-stranded RNA (ssRNA) viruses, produce dsRNA during their replication cycles [1–3]. Thus, cells have evolved a variety of dsRNA sensors which act as essential safeguards that detect and restrict viral invaders. Three important cytosolic dsRNA sensors widely found in human cells include RIG-I-like receptors (RLRs), 2’-5’ oligoadenylate synthetase (OAS), and protein kinase R (PKR) (Fig. 1A).

**Figure 1.**
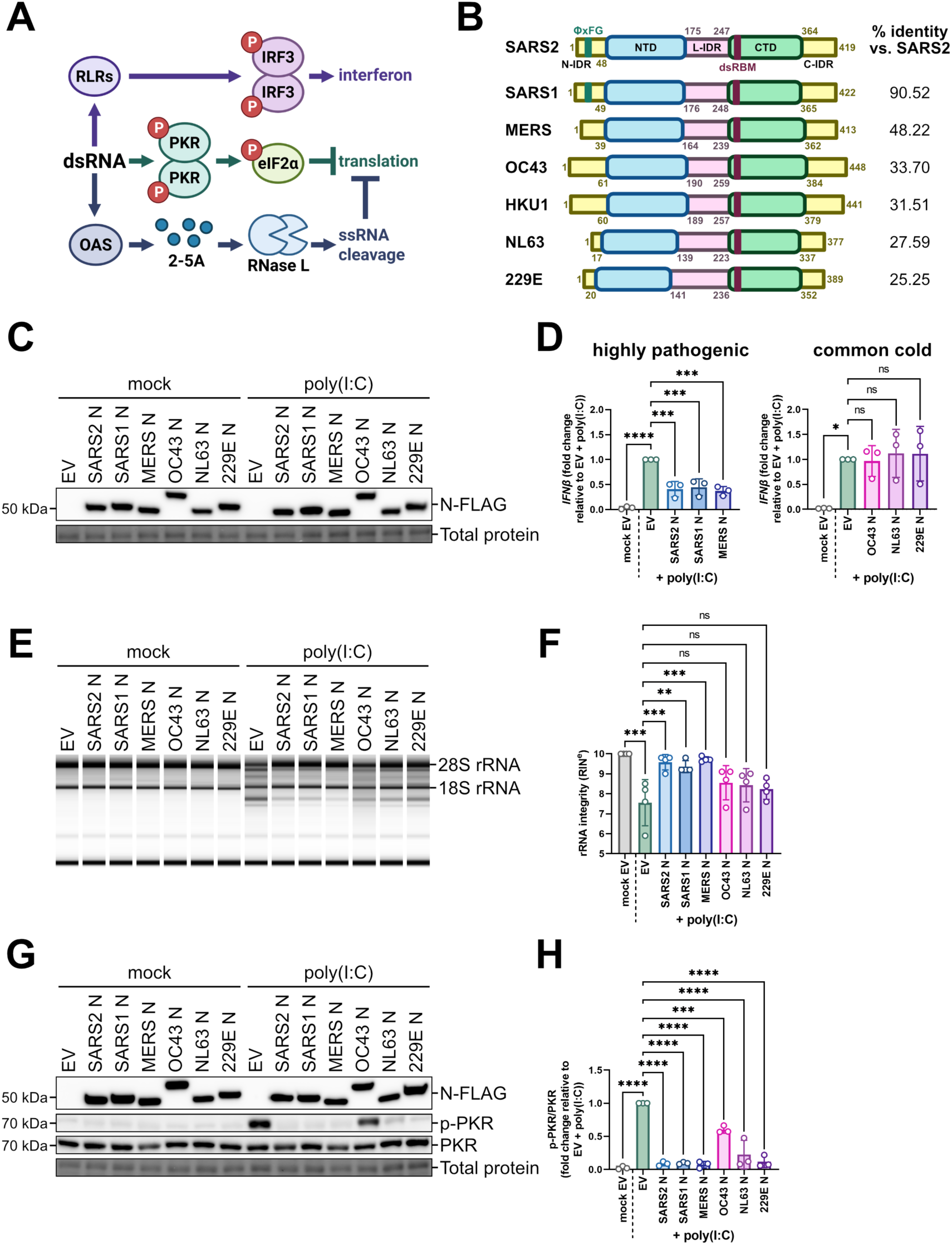
HCoV N proteins differentially inhibit the interferon, OAS/RNase L, and PKR pathways. **A.** Schematic depicting antiviral pathways induced by dsRNA. **B.** Schematic depicting the domain organization of human coronavirus N proteins and their percent identity (%) relative to SARS-CoV-2 N as determined by sequence alignment. **C.** A549 cells were transduced with recombinant lentiviruses expressing an empty vector control (EV) or each of the indicated HCoV N-FLAG constructs and selected with puromycin. For this experiment and all subsequent experiments, internal methionine residues in SARS-CoV-2 N constructs were mutated to ensure that only the full-length N proteoform was expressed (SARS2 N; M210I and M234V). Six days post-transduction, cells were mock-transfected or transfected with 0.5 µg poly(I:C). Protein lysates were harvested three hours post-poly(I:C) addition, resolved by SDS-PAGE, and subjected to immunoblotting with anti-FLAG antibody to detect N proteins. **D.** A549 cells were transduced, selected, and transfected with poly (I:C) as in C. Intracellular RNA was harvested three hours post-poly(I:C) addition and subjected to RT-qPCR to measure *IFNβ* mRNA. Values were normalized to 18S rRNA (housekeeping control) and are represented as fold change relative to the EV-transduced, poly(I:C)-transfected condition from three independent biological replicates (*n* = 3; mean ± SD). Statistics were performed using a one-way ANOVA with Dunnett’s post-hoc analysis (****, p < 0.0001; ***, p < 0.001; *, p < 0.05; ns, nonsignificant). **E.** A549 cells were transduced, selected, and transfected with poly(I:C) as in C. Intracellular RNA was harvested three hours post-poly(I:C) addition and subjected to automated electrophoresis using an Agilent 4200 TapeStation system to assess rRNA integrity. The electronic gel shown represents one of four independent biological replicates (*n* = 4). **F.** The RNA integrity number equivalent (RIN^e^) was measured for the samples analyzed by automated electrophoresis in E. These data represent three independent biological replicates (*n* = 3; mean ± SD). Statistics were performed using a one-way ANOVA with Dunnett’s post-hoc analysis (***, p < 0.001; **, p < 0.01; ns, nonsignificant).

RLRs include retinoic acid-inducible gene I (RIG-I) and melanoma differentiation-associated gene 5 (MDA5). Upon binding their dsRNA ligands, RLRs become activated, initiating a signaling cascade that results in the transcription of type I interferons (IFNs) including *IFNα* and *IFNβ* [4]. IFN proteins are then secreted and act in an autocrine or paracrine manner, resulting in the transcriptional activation of numerous interferon-stimulated genes (ISGs) with antiviral functions. The dsRNA sensors OAS and PKR are examples of ISGs, though they are also constitutively expressed in small amounts to allow cells to mount a rapid antiviral response [4]. Upon binding dsRNA, OAS synthesizes 2’-5’-oligoadenylates (2-5A), activating the endoribonuclease RNase L, which cleaves cytosolic host and viral ssRNAs. Meanwhile, PKR dimerizes when it binds dsRNA, resulting in its autophosphorylation and activation [1]. PKR is one of four integrated stress response (ISR) kinases which upon activation phosphorylate the translation initiation factor eIF2 on its alpha subunit (eIF2α), inducing translational arrest [5].

In addition to these canonical pathways, a growing body of work suggests that biomolecular condensates play important roles in the dsRNA-induced antiviral response. Translational arrest resulting from ISR activation leads to the formation of stress granules (SGs), large cytoplasmic condensates containing translationally arrested RNAs along with RNA-binding proteins such as the SG-nucleating protein G3BP1 [5,6]. SGs are thought to have antiviral functions, and accordingly their formation is inhibited by a variety of viruses [5,7–13]. G3BP1 and other RNA-binding proteins can also phase separate with cleaved RNAs produced by RNase L activation, forming smaller condensates called RNase L-induced bodies (RLBs) which may also have antiviral functions [14,15]. Biomolecular condensation is also thought to play a direct role in dsRNA sensing. Cytosolic dsRNAs produced during viral infection have recently been shown to form phase-separated condensates called dsRNA-induced foci (dRIFs), which act as hubs for innate immune signaling by recruiting and concentrating proteins such as PKR, OAS, and RNase L [16–21]. RIG-I has also recently been shown to form condensates in response to dsRNA that promote IFN signaling [22].

Coronaviruses are a diverse family of enveloped RNA viruses that cause a wide spectrum of disease severity when they infect humans. Among the seven human coronaviruses (HCoVs), the betacoronaviruses SARS-CoV-2 (the causative agent of COVID-19), SARS-CoV-1, and MERS-CoV, are termed “highly pathogenic” due to their ability to cause severe or fatal respiratory disease or other complications [23,24]. Meanwhile, the betacoronaviruses HCoV-OC43 and HCoV-HKU1, along with the alphacoronaviruses HCoV-NL63 and HCoV-229E, are referred to as seasonal or “common cold” HCoVs as they typically cause mild upper respiratory symptoms. Coronaviruses produce abundant dsRNAs as intermediates of viral RNA synthesis [2,3]; as such, to replicate successfully, they must evade dsRNA responses. The strategies used by highly pathogenic HCoVs to evade dsRNA responses have been a subject of extensive study [16,25–39]. However, the strategies used by common cold HCoVs to antagonize dsRNA responses remain poorly characterized, even though common cold HCoVs are widely used as models for studying HCoV biology under biosafety level 2 (BSL-2) conditions. Furthermore, while multiple HCoV structural, nonstructural, and accessory proteins have been implicated in innate immune evasion, the precise mechanisms by which these proteins inhibit dsRNA responses require further characterization. Here, we use a comparative approach to investigate how the essential viral nucleocapsid (N) protein antagonizes dsRNA responses during coronavirus infection.

The N protein is a multifunctional structural protein with essential roles in coronavirus packaging and viral RNA synthesis [40–44]. N proteins share a low degree of sequence similarity but have a conserved modular structure comprised of an amino-terminal globular domain (NTD) and carboxy-terminal globular domain (CTD), which are linked by an intrinsically-disordered linker (L-IDR) and flanked by amino- and carboxy-terminal intrinsically disordered regions (N- and C-IDR, respectively) (Fig. 1B) [43,45,46]. The NTD and CTD both have RNA-binding activity, but the NTD displays affinity for ssRNA or short, TRS-like sequences, while the CTD nonspecifically binds structured or dsRNA [47,48].

SARS-CoV-2 N inhibits the IFN, OAS/RNase L, and PKR pathways, and prevents the formation of SGs [9,26,35,38,39,49–57]. These functions have been attributed in part to a dsRNA-binding motif (dsRBM) within the SARS-CoV-2 N CTD, which is thought to sequester dsRNA from innate immune sensors [49,58–60]. Furthermore, some SARS-CoV-2 variants of concern including Omicron evolved to enhance production of an amino-terminally-truncated form of N, called N*^M210^ (hereafter referred to as N*) that is superior to full-length N at innate immune evasion and dsRNA binding and provides the virus with a fitness advantage [53,61–63]. The SARS-CoV-1 and MERS-CoV N proteins have also been shown to antagonize IFN and inhibit SGs [25,31,57], raising the possibility that evasion of dsRNA responses is a conserved function of HCoV N proteins. Indeed, all HCoV N proteins contain a dsRBM within their CTDs (Fig. 1B). However, no study has systematically compared the abilities of different HCoV N proteins to evade the IFN, OAS/RNase L, and PKR pathways, and the innate immune-evasive functions of common cold HCoV N proteins remain almost entirely uncharacterized. Moreover, the precise molecular processes underlying N-mediated evasion of dsRNA responses have not been fully elucidated.

Using a comparative approach to comprehensively define the abilities of HCoV N proteins to evade dsRNA pathways, we discover that innate immune evasion ability is not uniform across N proteins. While highly pathogenic HCoV N proteins antagonize the IFN and OAS/RNase L pathways, common cold HCoV N proteins do not. Although most HCoV N proteins block PKR activation and colocalize with dsRNA foci, HCoV-OC43 N does not, suggesting that differences in PKR inhibition ability are related to the ability of N proteins to interact with dsRNA. Contrary to prevailing assumptions, HCoV N proteins do not outcompete PKR for colocalization with dsRNA; instead, N proteins colocalize with PKR and dsRNA within dRIFs and prevent PKR activation in dRIFs. Finally, we show that HCoV N proteins differentially modulate the morphology and composition of G3BP1 foci. Together, these findings shed light on key mechanistic commonalities and differences in N innate immune evasion function across diverse coronaviruses.

## RESULTS

### HCoV N proteins differentially inhibit dsRNA responses

We sought to compare the ability of HCoV N proteins to evade dsRNA responses. To do this, we transduced A549 cells with recombinant lentiviruses expressing an empty vector (EV) control or the carboxy-terminally FLAG-tagged N proteins of the highly pathogenic HCoVs SARS-CoV-2, SARS-CoV-1, and MERS-CoV or the common cold HCoVs HCoV-OC43, HCoV-NL63, and HCoV-229E. We then transfected the cells with the synthetic dsRNA poly(I:C). SARS-CoV-2 N overexpression has previously been shown to produce truncated proteoforms N*^M210^ and N*^M234^ via leaky scanning [53]. To ensure that only full-length N was being expressed, for all overexpression experiments, we used a full-length SARS-CoV-2 N construct in which internal methionine codons 210 and 234 have been substituted with isoleucine and valine (M210I, M234V), respectively. We were able to obtain comparable levels of expression for these HCoV N proteins and we observed that N expression levels were unaltered by poly(I:C) transfection (Fig. 1C). We could not rescue a lentivirus expressing HCoV-HKU1 N, nor were we able to express HCoV-HKU1 N by transient transfection; thus, we excluded this N protein from our analyses.

To investigate the effect of HCoV N proteins on the IFN pathway, we measured *IFNβ* mRNA levels by RT-qPCR. As expected, *IFNβ* was strongly induced by poly(I:C), and expression of SARS-CoV-2 N, as well as SARS-CoV-1 and MERS-CoV N, decreased *IFNβ* levels approximately twofold (Fig. 1D). In contrast, HCoV N proteins from common cold coronaviruses did not inhibit *IFNβ* induction. Next, we evaluated the effect of HCoV N proteins on the OAS/RNase L pathway by using rRNA integrity as a proxy for RNase L activation. As expected, poly(I:C) transfection led to a significant decrease in rRNA integrity that was rescued by overexpression of SARS-CoV-2 N, indicating that SARS-CoV-2 N prevented poly(I:C)-induced RNase L activation (Fig. 1E-F). SARS-CoV-1 and MERS-CoV N also inhibited RNase L. By contrast, overexpression of the N proteins of common cold HCoVs only marginally restored rRNA integrity, and this increase was not statistically significant. Overall, these results suggest that the ability to inhibit IFN and OAS/RNase L is conserved across highly pathogenic HCoV N proteins, but that common cold HCoV N proteins are unable to block these pathways.

Finally, we investigated the ability of HCoV N proteins to inhibit the PKR pathway by immunoblotting for phospho-PKR. As expected, poly(I:C) transfection strongly induced PKR phosphorylation, and PKR activation was abrogated in the presence of SARS-CoV-2 N (Fig. 1G-H). PKR activation was also completely abrogated by overexpression of SARS-CoV-1, MERS-CoV, HCoV-NL63, and HCoV-229E N proteins. While HCoV-OC43 N also significantly inhibited PKR phosphorylation, this decrease in phospho-PKR was weaker than that observed for the other N proteins. Together, these results show that the ability to antagonize dsRNA responses is not conserved across all HCoV N proteins. Highly pathogenic HCoV N proteins antagonize IFN and OAS/RNase L, while common cold HCoVs are unable to do so. Furthermore, all N proteins inhibit PKR, but HCoV-OC43 N does so to a much lesser extent than other N proteins.

### Infection with two common cold HCoVs induces different antiviral responses

We wondered whether HCoV infection influences dsRNA responses in a manner that reflects our N overexpression experiments. To do this, we investigated the activation kinetics of the IFN, OAS/RNase L, and PKR pathways during infection using HCoV-OC43 as a representative betacoronavirus, and HCoV-229E as a representative alphacoronavirus. First, we sought to identify a cell type in which viral RNA accumulates to comparable levels during infection. Both viruses replicated to a high titre in A549 cells (Fig. S2A). However, we did not observe robust accumulation of viral genomes in infected A549 cells, especially in the case of HCoV-229E, where viral gRNA levels late in infection only increased approximately 40-fold relative to input gRNA (Fig. S2B-C). We predicted that this system would be problematic for assessing dsRNA responses during HCoV infection due to limited accumulation of immunogenic viral RNAs. Thus, we performed all further infection experiments in EA.Hy926 cells, an A549/human umbilical vein endothelial cell (HUVEC) hybrid cell line [64] which was permissive to both HCoV-OC43 and HCoV-229E (Fig. S2D). In contrast to what was observed during infection in A549 cells, both viruses exhibited robust gRNA replication in EA.Hy926 cells (Fig. S2E-F). Viral gRNA levels plateaued by approximately 24 hours post-infection (hpi) for HCoV-OC43 and between 12 and 24 hpi for HCoV-229E, suggesting that the viral replication cycle progresses faster for HCoV-229E than for HCoV-OC43.

We first assessed activation of the IFN pathway in our infection model by measuring levels of *IFNβ* mRNA as well as the ISG mRNAs *MX1, OAS1,* and *ISG15*. Neither HCoV-OC43 nor HCoV-229E induced detectable expression of *IFNβ* during infection (Fig. 2A-B). Both viruses had variable effects on ISG induction. *MX1* expression was induced late in infection by both viruses, with *MX1* levels increasing approximately threefold during HCoV-OC43 infection and fourfold during HCoV-229E infection (Fig. S3A, D). HCoV-229E infection also led to an increase in *OAS1* levels by 24 hpi (Fig. S3E). By contrast, HCoV-OC43 infection led to a strong reduction in *OAS1* expression by 12 hpi (Fig. S3B). Neither HCoV-OC43 nor HCoV-229E induced detectable expression of *ISG15* (Fig. S3C, F). Together, these results demonstrate that common cold coronaviruses induce some, but not all, ISGs during infection in EA.Hy926 cells, and that this induction may occur in an IFNβ-independent manner. Next, we investigated induction of the OAS/RNase L pathway during HCoV-OC43 and HCoV-229E infection. We did not observe rRNA cleavage during HCoV-OC43 or HCoV-229E infection (Fig. 2C-F), suggesting that neither of these viruses activate OAS/RNase L.

**Figure 2.**
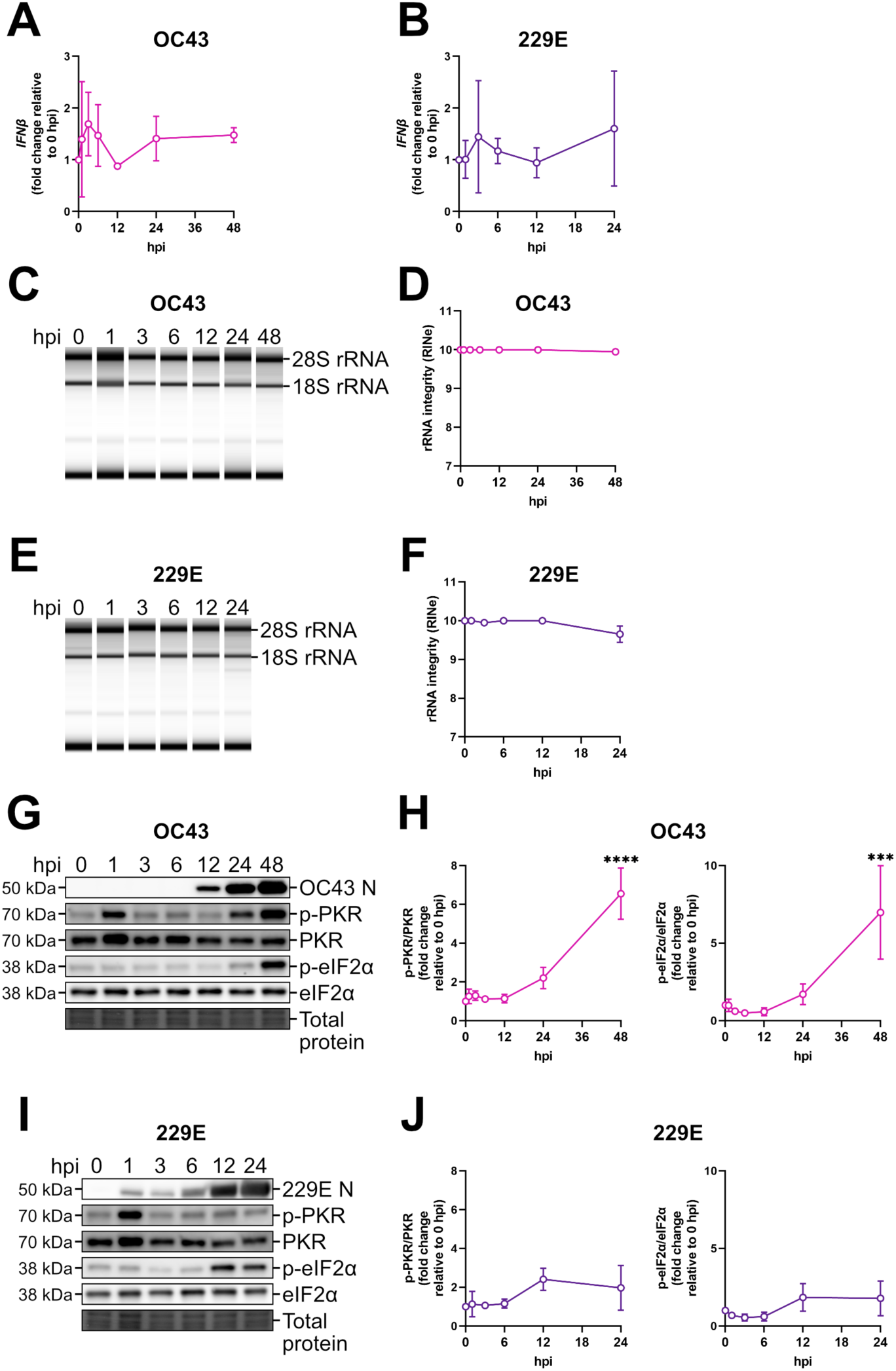
HCoV-OC43 and HCoV-229E differentially activate dsRNA-induced antiviral pathways during infection. **A-B.** EA.Hy926 cells were infected with HCoV-OC43 (A) or HCoV-229E (B) at an MOI of 1. Intracellular RNA was harvested at indicated times post-infection and subjected to RT-qPCR to measure *IFNβ* mRNA abundance. Values were normalized to 18S rRNA (housekeeping control) and are represented as fold change relative to 0 hpi from three independent biological replicates (*n* = 3; mean ± SD). Statistics were performed relative to 0 hpi using a one-way ANOVA with Dunnett’s post-hoc analysis (no symbol shown, nonsignificant). **C-F.** RNA from cells infected with HCoV-OC43 as in A. (C-D) or HCoV-229E as in B. (E-F) was subjected to automated electrophoresis using an Agilent 4200 TapeStation system to assess rRNA integrity. Representative electronic gels are shown (C, E). RNA integrity number equivalent (RIN^e^) values were measured for each sample (D, F). These data represent two independent biological replicates (*n* = 2; mean ± SD). **G-J.** EA.Hy926 cells were infected with HCoV-OC43 (G-H) or HCoV-229E (I-J) as in A-B. Protein lysates were harvested at indicated times post-infection, resolved by SDS-PAGE, and subjected to immunoblotting with antibodies specific to OC43 N (G), 229E N (I), p-PKR, PKR, p-eIF2α, and eIF2α. Representative immunoblots are shown (G, I). Protein levels were quantified by densitometry in ImageLab and normalized to total protein (H, J). Ratios of p-PKR to total PKR (left) and p-eIF2α to total eIF2α (right) were then calculated and normalized to 0 hpi. These data represent three independent biological replicates (*n* = 3; mean ± SD). Statistics were performed relative to 0 hpi using a one-way ANOVA with Dunnett’s post-hoc analysis ((****, p < 0.0001; ***, p < 0.001; no symbol shown, nonsignificant).

Finally, we interrogated induction of the PKR pathway during HCoV-OC43 and HCoV-229E infection. During infection with both viruses, we observed a transient increase in phospho-PKR levels at 1 hpi that was concomitant with an increase in total PKR, likely due to cell stress caused by the infection protocol (Fig. 2G, I). For both viruses, phospho-PKR levels returned to baseline by 3 hpi, but at late time points, we observed a marked difference in PKR induction between the two viruses. During HCoV-OC43 infection, there was an approximately 6.5-fold increase in mean phospho-PKR levels by 48 hpi, while during HCoV-229E infection, there was no significant increase in phospho-PKR (Fig. 2G-J). We also evaluated phospho-eIF2α levels during infection and observed that HCoV-OC43 induced a sevenfold increase in eIF2α phosphorylation by 48 hpi (Fig. 2G-H). Meanwhile, HCoV-229E induced an approximately 1.8-fold increase in eIF2α phosphorylation by 12 hpi that was not statistically significant (Fig. 2I-J). Together, these data show that HCoV-OC43 activates the PKR pathway to a much greater extent than HCoV-229E; this activation of PKR likely contributes to increased eIF2α phosphorylation, though other ISR kinases such as PERK likely also play a role in phosphorylating eIF2α during HCoV-OC43 infection [65,66]. Furthermore, while both viruses induce eIF2α phosphorylation, HCoV-229E-mediated induction of eIF2α is weaker and likely occurs via ISR kinases other than PKR [67,68].

### Common cold HCoVs differ in their ability to antagonize antiviral responses induced by exogenous dsRNA

Our initial infection experiments suggested that both HCoV-OC43 and HCoV-229E antagonize *IFNβ* induction and OAS/RNase L activation, and that HCoV-229E, but not HCoV-OC43 inhibits PKR activation. However, one alternative explanation for a lack of activation of these pathways during infection is that one or both viruses may not produce sufficient cytosolic dsRNA to elicit an innate immune response. Another alternative explanation is that EA.Hy926 cells are unable to mount an immune response to coronavirus RNA; for instance, these cells may lack the correct OAS1/2 protein isoforms required to detect membrane-bound dsRNA intermediates generated during coronavirus infection [69–72]. To rule out these possibilities, we investigated whether HCoV-OC43 and HCoV-229E can antagonize dsRNA responses induced by a large amount of exogenous dsRNA. We infected EA.Hy926 cells with HCoV-OC43 or HCoV-229E, then transfected the cells with poly(I:C) and assessed activation of innate immune pathways (Fig. 3A).

**Figure 3.**
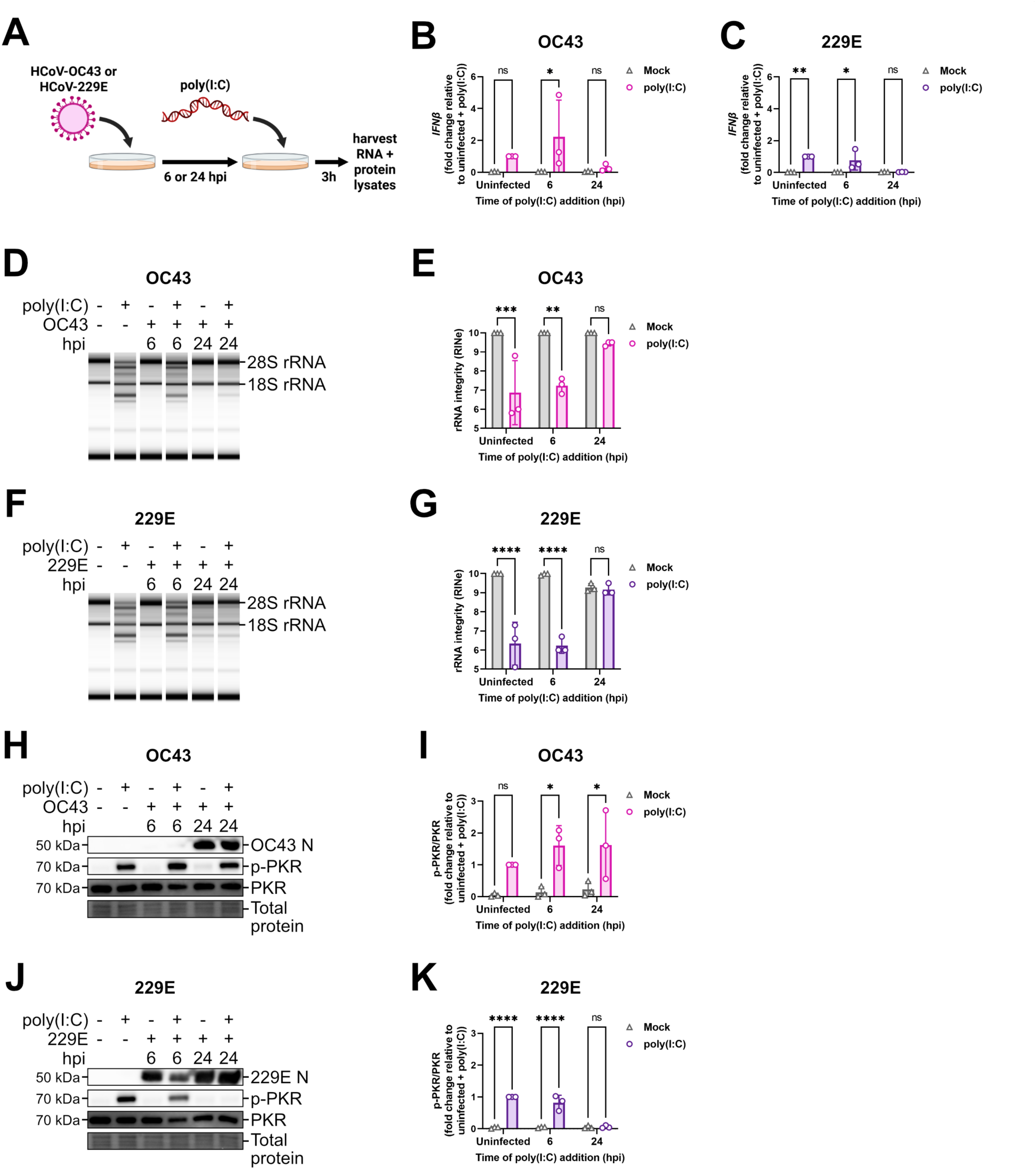
HCoV-OC43 and HCoV-229E differentially antagonize antiviral responses induced by exogenous dsRNA. **A.** Schematic depicting protocol for assessing the effect of HCoV infection on antiviral responses induced by exogenous dsRNA. **B-C.** EA.Hy926 cells were mock-infected or infected with HCoV-OC43 (B) or HCoV-229E (C) at an MOI of 1, then mock-transfected or transfected with 0.5 µg poly(I:C) at the indicated time points post-infection. Intracellular RNA was harvested three hours post-poly(I:C) addition and subjected to RT-qPCR to measure *IFNβ* mRNA abundance. Values were normalized to 18S rRNA (housekeeping control) and are represented as fold change relative to the mock-infected, poly(I:C)-transfected condition from three independent biological replicates (*n* = 3; mean ± SD). Statistics were performed using a two-way ANOVA (**, p < 0.01; *, p < 0.05; ns, nonsignificant). **D-G.** RNA from B-C. was subjected to automated electrophoresis using an Agilent 4200 TapeStation system to assess rRNA integrity. Representative gels are shown (D, F). RNA integrity number equivalent (RIN^e^) values were measured for each sample (E, G). These data represent three independent biological replicates (*n* = 3; mean ± SD). Statistics were performed using a two-way ANOVA (****, p < 0.0001; ***, p < 0.001; **, p < 0.01; ns, nonsignificant). **H-K.** EA.Hy926 cells were infected with HCoV-OC43 (H-I) or HCoV-229E (J-K) and transfected with poly(I:C) as in B-C. Protein lysates were harvested three hours post-poly(I:C) addition, resolved by SDS-PAGE, and subjected to immunoblotting with antibodies specific to OC43 N (H), 229E N (J), p-PKR, and PKR. Representative immunoblots are shown (H, J). Protein levels were quantified by densitometry in ImageLab and normalized to total protein (I, K). Ratios of p-PKR to total PKR were then calculated and normalized to the mock-infected, poly(I:C)-transfected condition. These data represent three independent biological replicates (*n* = 3, mean ± SD). Statistics were performed using a two-way ANOVA (****, p < 0.0001; *, p < 0.05; ns, nonsignificant).

In uninfected EA.Hy926 cells transfected with poly(I:C), we observed an increase in *IFNβ* levels, rRNA cleavage, and PKR phosphorylation, indicating the IFN, OAS/RNase L, and PKR pathways were robustly induced by poly(I:C) (Fig. 2). However, by 24 hpi, we observed that both HCoV-OC43 and HCoV-229E completely blocked *IFNβ* induction by poly(I:C) (Fig. 3B-C). Both viruses also inhibited RNase L activation by poly(I:C) at 24 hpi (Fig. 3D-G). However, the two viruses strongly differed in their ability to antagonize PKR; while HCoV-OC43 was unable to block poly(I:C)-induced PKR activation, HCoV-229E completely prevented PKR activation when cells were transfected with poly(I:C) 24 hpi (Fig. 3H-K). These results corroborate our findings from our initial infection experiments, confirming that HCoV-OC43 and HCoV-229E actively antagonize dsRNA-induced *IFNβ* induction and RNase L activation, and that HCoV-229E antagonizes PKR while HCoV-OC43 does not. From these findings, we have the following interpretations: i) N contributes to PKR inhibition during HCoV-229E infection, while HCoV-OC43 infection induces PKR activation because its N protein is a weaker PKR antagonist and ii) viral proteins other than N are likely responsible for inhibiting *IFNβ* and blocking OAS/RNase L during HCoV-OC43 and HCoV-229E infection.

### N proteins that inhibit PKR colocalize with dsRNA foci

We wondered how HCoV N proteins block the IFN, OAS/RNase L, and PKR pathways. Previous reports suggest that SARS-CoV-2 N inhibits innate immune responses via a dsRNA-binding motif (dsRBM) comprised of two lysines within its CTD [49,53,58]. Given that these two lysines are conserved across the N proteins of all HCoVs (Fig. 1B, Fig. S1, Fig. 4A), we wondered whether dsRNA interaction ability is a shared feature of HCoV N proteins. To examine this, we overexpressed each HCoV N protein in A549 cells, transfected the cells with poly(I:C), and immunostained for N and dsRNA to evaluate N enrichment within dsRNA foci as a proxy for N-dsRNA interaction [53]. As a positive control for N-dsRNA colocalization, we also overexpressed SARS-CoV-2 N*, because this truncated SARS-CoV-2 N proteoform has enhanced dsRNA interaction ability [53].

**Figure 4.**
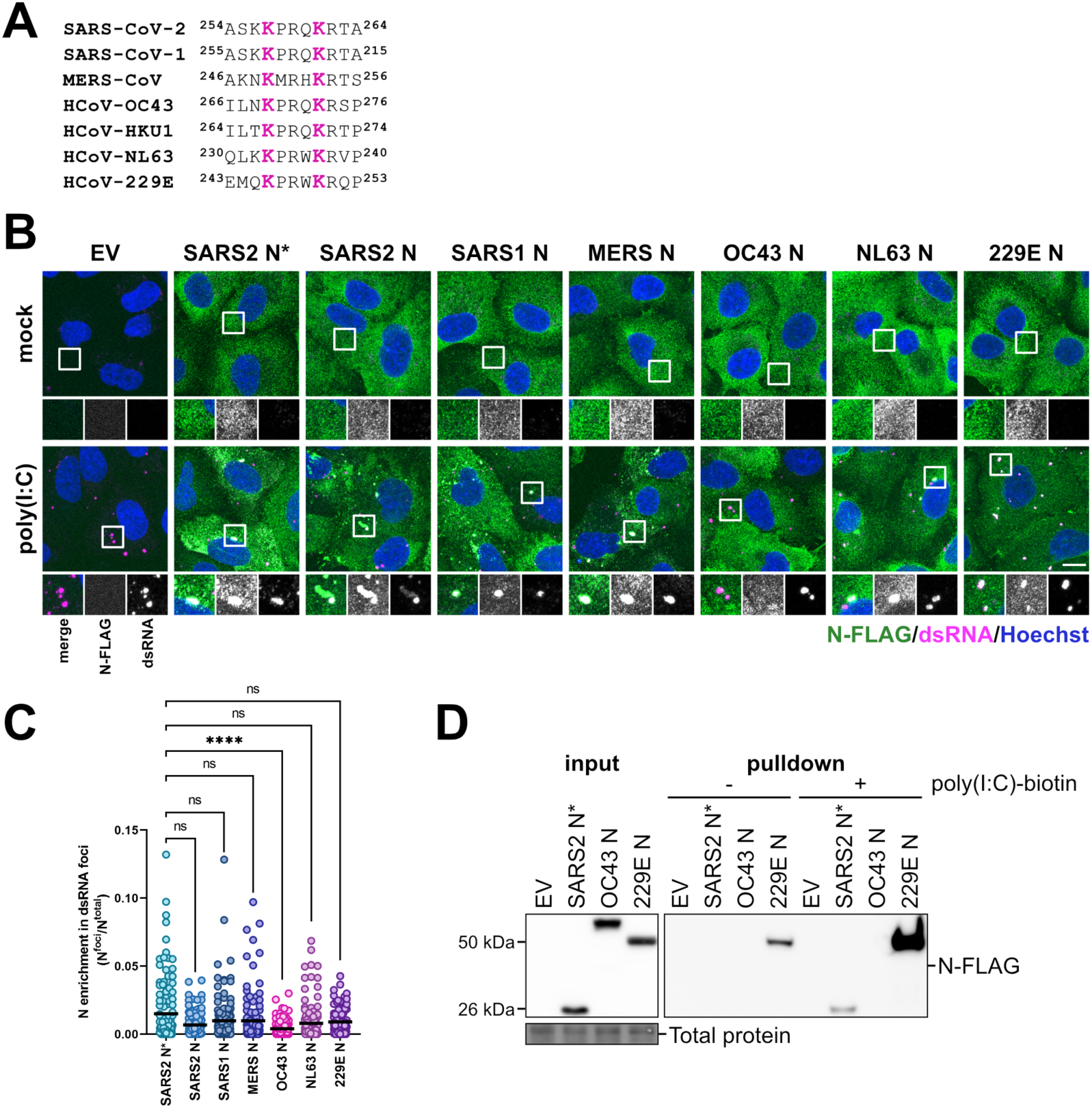
N proteins other than HCoV-OC43 N colocalize with dsRNA foci. **A.** Amino acid sequence surrounding the dsRNA-binding motif (dsRBM) (magenta) of HCoV N proteins. **B.** A549 cells were transduced with recombinant lentiviruses expressing an empty vector control (EV) or each of the indicated HCoV N-FLAG constructs and selected with puromycin. Six days post-transduction, cells were mock-transfected or transfected with 0.5 µg poly(I:C). Cells were fixed three hours post-poly(I:C) addition and immunostained with antibodies specific to FLAG (N protein; green) or dsRNA (magenta). Nuclei were stained with Hoechst (blue). Images were obtained using an LSM 880 Airyscan confocal microscope, with maximum-intensity projections presented (scale bar = 10 µm). **C.** The ratio of N intensity within dsRNA foci (N^foci^) relative to total N intensity (N^total^) per N-expressing cell from B. was quantified in CellProfiler. At least 25 cells were imaged per biological replicate (*n* = 3; mean). Statistics were performed using a Kruskal-Wallis H test with Dunn’s post-hoc analysis (****, p < 0.0001; ns, nonsignificant). **D.** Cells were transduced and selected as in B. Four days post-transduction, protein lysates were harvested and precipitated using streptavidin beads conjugated to biotinylated poly(I:C) or unconjugated beads. Eluted proteins were subjected to SDS-PAGE and immunoblotting with antibodies specific to FLAG (N proteins). The immunoblot shown represents one of three independent biological replicates (*n* = 3).

All HCoV N proteins displayed a diffuse localization in the absence of poly(I:C) (Fig. 4B). As previously reported, upon poly(I:C) transfection, SARS-CoV-2 N* became strongly enriched within dsRNA foci, while full-length SARS-CoV-2 N partially colocalized with dsRNA foci (Fig. 4B-C) [53]. Full-length SARS-CoV-2 N also localized to irregular-shaped puncta lacking dsRNA which have been previously identified as stress granules [53,73]. Like SARS-CoV-2 N, SARS-CoV-1 N was enriched in both round dsRNA-containing foci and irregularly-shaped, non-dsRNA-containing punta, likely also stress granules [74]. The N proteins of MERS-CoV, HCoV-NL63, and HCoV-229E also became enriched in dsRNA foci upon poly(I:C) transfection. By contrast to all other HCoV N proteins tested, HCoV-OC43 N failed to significantly colocalize with dsRNA foci and did not form other puncta, retaining its diffuse localization. These results demonstrate that most HCoV N proteins can colocalize with dsRNA to some extent, except for HCoV-OC43 N.

To validate that N-dsRNA colocalization phenotypes represent an N-dsRNA interaction, we collected lysates from A549 cells expressing HCoV-OC43 N, HCoV-229E N, or SARS-CoV-2 N* as a positive control and performed a pulldown assay using beads conjugated to biotinylated poly(I:C). As expected, SARS-CoV-2 N* was absent in the eluate obtained from unconjugated beads but was detected in the eluate obtained from poly(I:C)-conjugated beads, showing that SARS-CoV-2 N* interacts with dsRNA (Fig. 4D). While some HCoV-229E N was observed in the eluate obtained from unconjugated beads, this N protein was strongly enriched in the eluate from poly(I:C)-conjugated beads, while HCoV-OC43 N was not detected. This suggests that HCoV-229E N interacts with dsRNA while HCoV-OC43 N does not, corroborating the results of our immunofluorescence experiments.

Taken together with our previous data, these experiments reveal that some, but not all, N-mediated innate immune evasion phenotypes correlate with dsRNA binding ability (Table 1). HCoV-OC43 N was unable to interact with dsRNA (Fig. 4) and was unable to strongly antagonize PKR activation (Fig. 1), raising the possibility that N-dsRNA interaction may be important for blocking PKR. By contrast, while HCoV-NL63 and HCoV-229E N were able to colocalize with dsRNA, they were unable to block IFN or OAS/RNase L activation (Fig. 1), showing that dsRNA colocalization is not sufficient to enable N-mediated inhibition of IFN and OAS/RNase L.

**Table 1.** Summary of dsRNA response antagonism by HCoVs and their N proteins.

|  | HCoV N overexpression |  |  |  |  |  | HCoV infection |  |
| --- | --- | --- | --- | --- | --- | --- | --- | --- |
|  | SARS2 | SARS1 | MERS | OC43 | NL63 | 229E | OC43 | 229E |
| *IFN*β inhibition | + | + | + | - | - | - | ++ | ++ |
| RNase L inhibition | + | + | + | - | - | - | ++ | ++ |
| PKR inhibition | ++ | ++ | ++ | -/+ | ++ | ++ | - | ++ |
| Colocalization with dsRNA | + | ++ | ++ | - | ++ | ++ | ND | ND |

*ND = not determined.*

### The dsRNA binding motif is required for N-mediated inhibition of dsRNA responses

Prior studies have attributed SARS-CoV-2 N-mediated innate immune evasion to the dsRBM, which was shown to be necessary for dsRNA binding and inhibition of RLRs, OAS, and PKR; however, it is not known whether this motif is sufficient to confer these functions [49,53,59]. Our findings indicate that conservation of the dsRBM across HCoV N proteins (Fig. S1, Fig. 4A) is not sufficient for dsRNA binding and innate immune evasion, because HCoV-OC43 N failed to colocalize with dsRNA or effectively inhibit PKR; furthermore, neither HCoV-OC43 N nor HCoV-NL63 and HCoV-229E N were able to block IFN and OAS/RNase L. Thus, we wondered if the dsRBM is necessary for N-mediated innate immune evasion in our system. To investigate this, we selected full-length SARS-CoV-2 N and HCoV-229E N as representative highly pathogenic and common cold HCoV N proteins that contain a dsRBM and co-precipitate and colocalize with dsRNA. We then generated SARS-CoV-2 and HCoV-229E N constructs in which the dsRBM lysines were eliminated (SARS2 N^ΔRBM^; K257A/K261A and 229E N^ΔRBM^; K246R/K250R, respectively).

Using immunostaining, we verified the ability of each N construct to colocalize with dsRNA. SARS2 N^ΔRBM^ was diffusely cytoplasmic in the absence of poly(I:C), but unlike wild-type SARS-CoV-2 N, it failed to colocalize with dsRNA (Fig. S4). 229E N^ΔRBM^ formed large cytoplasmic puncta in the absence of poly(I:C), suggesting that the K246R/K250R double substitution caused protein aggregation. After poly(I:C) transfection, 229E N^ΔRBM^ puncta did not colocalize with dsRNA. Together, these observations confirm that the dsRBM of SARS-CoV-2 and HCoV-229E N is required for the proteins to colocalize with dsRNA.

Next, we evaluated the ability of the dsRBM mutant constructs to inhibit innate immune responses. Neither SARS2 N^ΔRBM^ nor 229E N^ΔRBM^ were able to reduce *IFNβ* levels or prevent rRNA cleavage, showing that they were incapable of inhibiting IFN and RNase L responses (Fig. 5A-C). SARS2 N^ΔRBM^ and 229E N^ΔRBM^ were both unable to block PKR activation (Fig. 5D-E). Overall, these results demonstrate that the dsRBM is necessary, though not sufficient, to confer N-mediated dsRNA binding and innate immune evasion ability.

**Figure 5.**
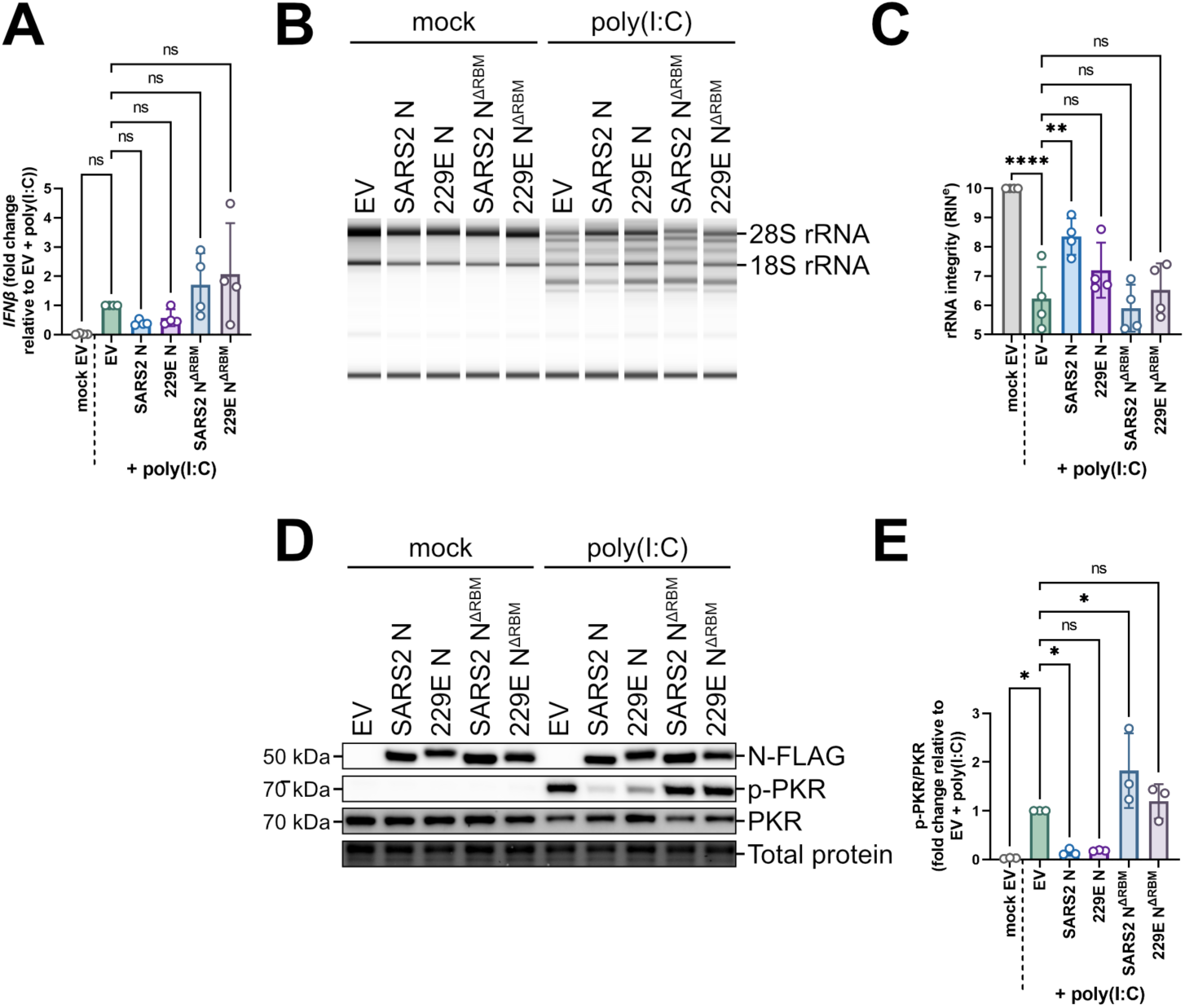
The dsRNA-binding motif (dsRBM) is necessary for N-mediated evasion of dsRNA responses. **A.** A549 cells were transduced with recombinant lentiviruses expressing an empty vector control (EV) or each of the indicated HCoV N-FLAG constructs and selected with puromycin. The dsRBM lysines were mutated to alanine for the SARS2^ΔRBM^ construct (K257A/K261A) or arginine for the 229E^ΔRBM^ construct (K246R/K250R). Six days post-transduction, cells were mock-transfected or transfected with 0.5 µg poly(I:C). Intracellular RNA was harvested three hours post-poly(I:C) addition and subjected to RT-qPCR to measure *IFNβ* mRNA abundance. Values were normalized to 18S rRNA (housekeeping control) and are represented as fold change relative to the EV-transduced, poly(I:C)-transfected condition from four independent biological replicates (*n* = 4; mean ± SD). Statistics were performed using a one-way ANOVA with Dunnett’s post-hoc analysis (ns, nonsignificant). **B.** A549 cells were transduced, selected, and transfected with poly(I:C) as in A. Intracellular RNA was harvested three hours post-poly(I:C) addition and subjected to automated electrophoresis using an Agilent 4200 TapeStation system to assess rRNA integrity. The electronic gel shown represents one of three independent biological replicates (*n* = 3). **C.** The RNA integrity number equivalent (RIN^e^) was measured for the samples analyzed by automated electrophoresis in B. These data represent three independent biological replicates (*n* = 3; mean ± SD). Statistics were performed using a one-way ANOVA with Dunnett’s post-hoc analysis ((****, p < 0.0001; **, p < 0.01; ns, nonsignificant). **D.** A549 cells were transduced, selected, and transfected with poly(I:C) as in A. Protein lysates were harvested three hours post-poly(I:C) addition, resolved by SDS-PAGE, and subjected to immunoblotting with antibodies specific to FLAG (N proteins), p-PKR, and PKR. The immunoblot shown represents one of three independent biological replicates (*n* = 3). **E.** Protein levels from D. were quantified by densitometry in ImageLab and normalized to total protein. Ratios of p-PKR to total PKR were then calculated and normalized to the EV-transduced, poly(I:C)-transfected condition. These data represent three independent biological replicates (*n* = 3; mean ± SD). Statistics were performed using a one-way ANOVA with Dunnett’s post-hoc analysis (*, p < 0.05; ns, nonsignificant).

### The innate immune evasion abilities of N proteins are partially determined by their carboxy terminus

Based on our findings, other regions beyond the dsRBM are required for dsRNA colocalization and innate immune antagonism by HCoV N proteins. Given that the dsRBM is located within the CTD of N, we wondered whether additional features of the CTD and/or C-IDR were involved. To investigate this, we created chimeric mutants wherein the carboxy terminus of HCoV-OC43 N, the only N protein entirely unable to interact with dsRNA and inhibit dsRNA responses, was swapped with the carboxy termini of other N proteins: i) “OC-SARS2” contains the CTD and C-IDR of SARS-CoV-2 N, and ii) “OC-229E” contains the CTD and C-IDR of HCoV-229E N (Fig. 6A). We assessed the ability of these chimeric mutants to colocalize with dsRNA via immunofluorescence microscopy. In contrast to HCoV-OC43 N, which did not colocalize with dsRNA foci as expected, both OC-SARS2 and OC-229E chimeras were enriched in dsRNA foci (Fig. 6B-C), although this enrichment was greater for OC-SARS2 compared to OC-229E. These results suggest that the carboxy terminus is sufficient to enable dsRNA colocalization for SARS-CoV-2 N; however, HCoV-229E N utilizes other motifs to confer maximal dsRNA colocalization ability.

**Figure 6.**
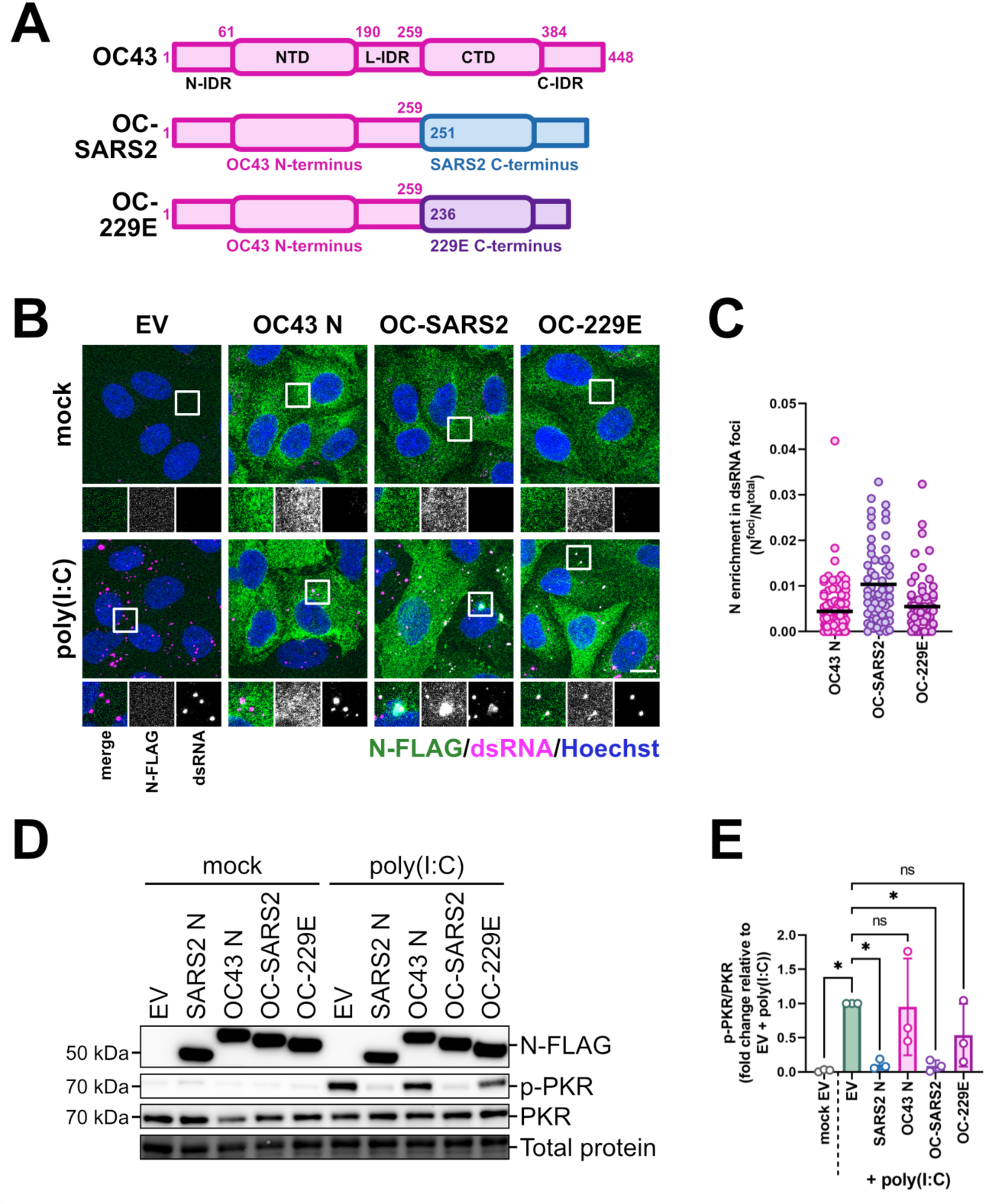
The carboxy termini of HCoV N proteins contribute to their ability to co-localize with dsRNA and block PKR activation. **A.** Schematic depicting chimeric HCoV-OC43 N mutants. In the OC-SARS2 mutant, the CTD and C-IDR of HCoV-OC43 were swapped with the CTD and C-IDR (residue 251 and onward) of SARS-CoV-2 N. In the OC-229E mutant, the CTD and C-IDR of HCoV-OC43 N were swapped with the CTD and C-IDR (residue 236 and onward) of HCoV-229E N. **B.** A549 cells were transduced with recombinant lentiviruses expressing an empty vector control (EV) or each of the indicated HCoV N-FLAG constructs and selected with puromycin. Six days post-transduction, cells were mock-transfected or transfected with 0.5 µg poly(I:C). Cells were fixed three hours post-poly(I:C) addition and immunostained with antibodies specific to FLAG (N protein; green) or dsRNA (magenta). Nuclei were stained with Hoechst (blue). Images were obtained using an LSM 880 Airyscan confocal microscope, with maximum-intensity projections presented (scale bar = 10 µm). The images shown represent one of two independent biological replicates (*n* = 2). **C.** The ratio of N intensity within dsRNA foci (N^foci^) relative to total N intensity (N^total^) per N-expressing cell was quantified in CellProfiler. At least 25 cells were imaged per biological replicate (*n* = 2; mean). **D.** A549 cells were transduced, selected, and transfected with poly(I:C) as in B. Protein lysates were harvested three hours post-poly(I:C) addition, resolved by SDS-PAGE, and subjected to immunoblotting with antibodies specific to FLAG (N proteins), p-PKR, and PKR. The immunoblot shown represents one of three independent biological replicates (*n* = 3). **E.** Protein levels from D. were quantified by densitometry in ImageLab and normalized to total protein. Ratios of p-PKR to total PKR were then calculated and normalized to the EV-transduced, poly(I:C)-transfected condition. These data represent three independent biological replicates (*n* = 3; mean ± SD). Statistics were performed using a one-way ANOVA with Dunnett’s post-hoc analysis (*, p < 0.05; ns, nonsignificant).

Next, we characterized the ability of the chimeric mutants to inhibit dsRNA responses. While HCoV-OC43 N did not inhibit *IFNβ* induction by poly(I:C), both OC-SARS2 and OC-229E chimeras inhibited *IFNβ* induction to the same extent as wild-type, full-length SARS-CoV-2 N (Fig. S5A). Moreover, both chimeras reduced poly(I:C)-induced rRNA cleavage to the same extent as SARS-CoV-2 N, showing that they were able to block RNase L activation (Fig. S5B-C). By contrast, only the OC-SARS2 mutant was able to inhibit PKR to the same extent as wild-type SARS-CoV-2 N, while expression of the OC-229E mutant led to a modest decrease in phospho-PKR levels that was not statistically significant (Fig. 6D-E).

These observations enabled us to draw four main inferences about how individual domains of N influence protein function. First, dsRNA colocalization ability is partially conferred by the CTD and/or C-IDR, but in some cases, like HCoV-229E N, additional regions are required for optimal dsRNA interaction. Second, dsRNA colocalization ability of N correlates with the ability to inhibit PKR. Third, the ability of N to block IFN and OAS/RNase L is independent from its ability to colocalize with dsRNA or inhibit PKR. Fourth, IFN and RNase L inhibition ability is likely determined by both the amino and carboxy termini of HCoV N proteins, because both the OC-SARS2 and OC-229E chimeras blocked IFN and RNase L, even though wild-type HCoV-229E N did not inhibit these pathways.

### N proteins that interact with dsRNA co-condense with PKR within dsRNA-induced foci (dRIFs)

PKR activation has been shown to occur within dsRNA- and PKR-containing condensates called dsRNA-induced foci (dRIFs), which may enhance PKR activation by concentrating PKR and dsRNA [18]. A recent study demonstrated that MERS-CoV dsRNA binding protein NS4a outcompetes PKR for condensation on dsRNA, preventing dRIF assembly and blocking PKR activation [16]. Because the ability of N proteins to decrease PKR phosphorylation and their ability to colocalize with dsRNA correlate, we hypothesized that N proteins may inhibit PKR by competing for dsRNA binding, blocking dRIF formation.

A549 cells expressing each N protein were transfected with poly(I:C), then immunostained for N, dsRNA, and PKR. In control cells, PKR colocalized with dsRNA, a marker of dRIF formation (Fig. 7A). N proteins other than HCoV-OC43 N colocalized with dsRNA foci, as in Fig. 4B-C. Contrary to our expectation, the percentage of cells containing PKR foci remained unchanged by N overexpression, showing that N proteins do not inhibit dRIF formation (Fig. 7B). Furthermore, dsRNA foci containing N proteins did not exclude PKR; rather, N and PKR both colocalized with dsRNA, a finding that is not consistent with a competitive model (Fig. 7A). We also observed that while MERS-CoV, HCoV-NL63, and HCoV-229E N proteins were enriched within large, round PKR foci that typically contained dsRNA, SARS-CoV-2 and SARS-CoV-1 N were enriched in dsRNA-containing PKR foci as well as within irregular PKR condensates which did not contain dsRNA and resembled stress granules (Fig. 7A, C). HCoV-OC43 N was the only N protein that was not enriched within either type of PKR-positive foci (Fig. 7A, C).

**Figure 7.**
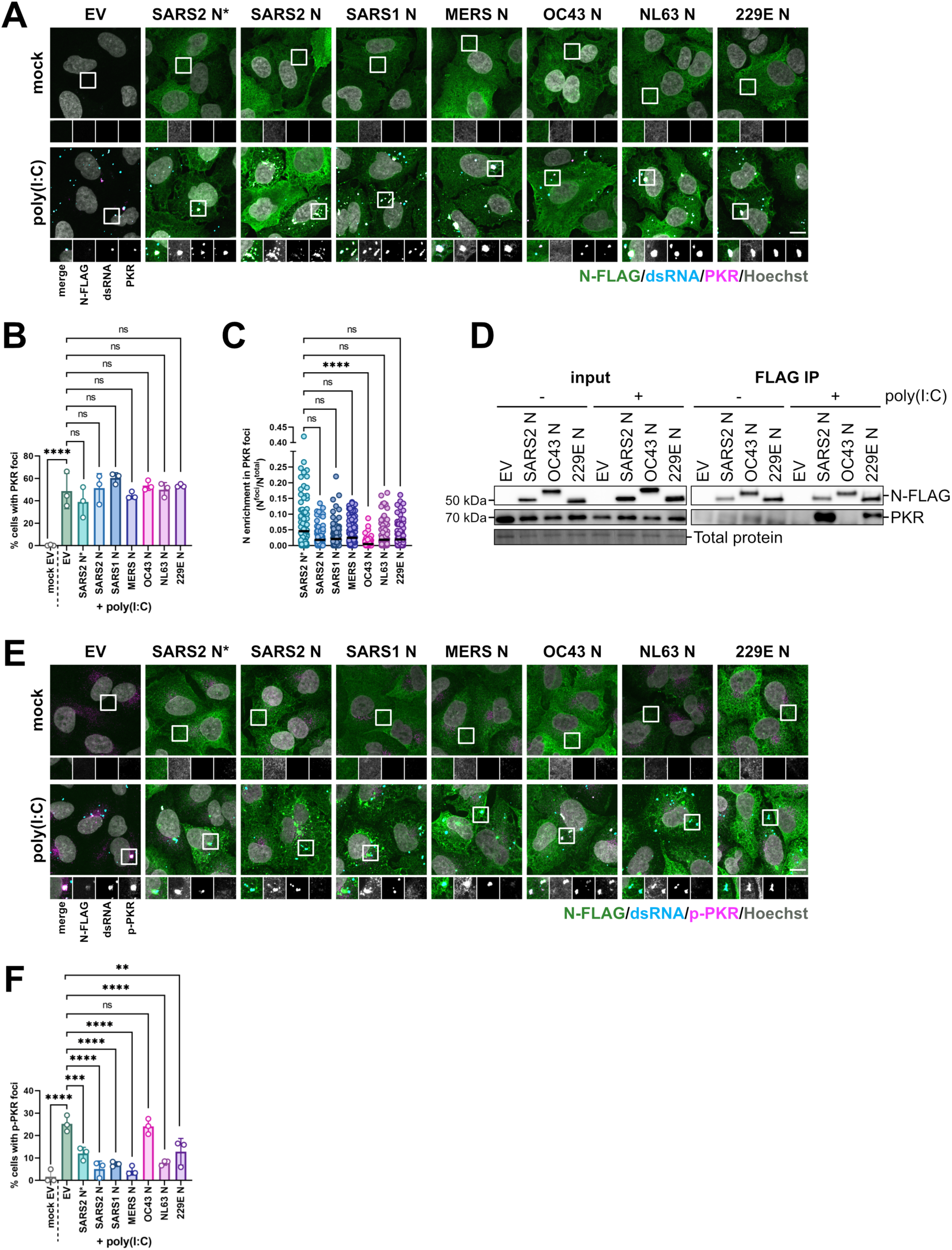
N proteins except HCoV-OC43 N colocalize with PKR and dsRNA and block PKR activation. **A.** A549 cells were transduced with recombinant lentiviruses expressing an empty vector control (EV) or each of the indicated HCoV N-FLAG constructs and selected with puromycin. Six days post-transduction, cells were mock-transfected or transfected with 0.5 µg poly(I:C). Cells were fixed three hours post-poly(I:C) addition and immunostained with antibodies specific to FLAG (N protein; green), dsRNA (cyan), or PKR (magenta). Nuclei were stained with Hoechst (grey). Images were obtained using an LSM 880 Airyscan confocal microscope, with maximum-intensity projections presented (scale bar = 10 µm). **B.** Coverslips from A. were imaged using a Zeiss AxioObserver Z1 microscope. The percentage of total cells (for cells transduced with EV) or N-expressing cells (for cells transduced with HCoV N constructs) containing PKR foci was quantified in CellProfiler. At least 171 cells were imaged per biological replicate (*n* = 3; mean ± SD). Statistics were performed using a one-way ANOVA with Dunnett’s post-hoc analysis (****, p < 0.0001; ns, nonsignificant)… **C.** The ratio of N intensity within PKR foci (N^foci^) relative to total N intensity (N^total^) per N-expressing cell in confocal images from A. was quantified in CellProfiler. At least 37 cells were imaged per biological replicate (*n* = 3; mean). Statistics were performed using a Kruskal-Wallis H test with Dunn’s post-hoc analysis (****, p < 0.0001; ns, nonsignificant). **D.** A549 cells were transduced and selected as in A. Four days post-transduction, cells were mock-transfected or transfected with 0.5 µg poly(I:C). Three hours post-poly(I:C) addition, protein lysates were harvested and N proteins were immunoprecipitated using an antibody specific to FLAG. Eluted proteins were subjected to SDS-PAGE and immunoblotted using antibodies specific to FLAG (N proteins) and PKR. The immunoblot shown represents one of three independent biological replicates (*n* = 3). **E.** A549 cells were transduced, selected, and transfected with poly(I:C) as in A. Cells were fixed three hours post-poly(I:C) addition and immunostained with antibodies specific to FLAG (N protein; green), dsRNA (cyan), or p-PKR (magenta). Nuclei were stained with Hoechst (grey). Images were obtained using an LSM 880 Airyscan confocal microscope, with maximum-intensity projections presented (scale bar = 10 µm). **F.** Coverslips from E. were imaged using a Zeiss AxioObserver Z1 microscope. The percentage of total cells (for cells transduced with EV) or N-expressing cells (for cells transduced with HCoV N constructs) containing p-PKR foci was quantified in CellProfiler. At least 147 cells were imaged per biological replicate (*n* = 3; mean ± SD). Statistics were performed using a one-way ANOVA with Dunnett’s post-hoc analysis (****, p < 0.0001; ***, p < 0.001; **, p < 0.01; ns, nonsignificant).

Given that HCoV-OC43 N was the only N protein unable to interact with dsRNA, we wondered if N proteins co-precipitate with PKR in a dsRNA-dependent manner. We overexpressed a representative subset of FLAG-tagged N proteins, comprising SARS-CoV-2, HCoV-OC43, and HCoV-229E N, in A549 cells. We transfected the cells with poly(I:C) or a mock control, collected protein lysates, and performed co-immunoprecipitation using beads conjugated to an anti-FLAG antibody. Immunoblotting revealed that when cells were not transfected with poly(I:C), PKR did not co-precipitate with any N protein (Fig. 7D). However, when cells were transfected with poly(I:C), PKR was detected in the immunoprecipitated eluate from SARS-CoV-2 and HCoV-229E N-expressing cells, suggesting that dsRNA is required for N-PKR interaction. HCoV-OC43 N was unable to interact with PKR, even in the presence of poly(I:C). Taken together, these results show that HCoV N proteins do not inhibit dRIF formation and do not compete with PKR for access to dsRNA; instead, N proteins localize with PKR within dRIFs in a dsRNA-dependent manner.

### N proteins that interact with dsRNA interfere with PKR phosphorylation within dRIFs

If N proteins do not competitively sequester dsRNA from PKR, how do they inhibit PKR activation? We hypothesized that N-PKR colocalization may alter PKR phosphorylation within dRIFs. To test this, we immunostained for N, dsRNA, and phospho-PKR. Phospho-PKR was not detected in the absence of poly(I:C), while poly(I:C) transfection led to the formation of foci positive for phospho-PKR and dsRNA in control cells (Fig. 7E-F). Although poly(I:C) transfection induced PKR-containing dRIFs in approximately 50% of cells, only approximately 25% of cells contained phospho-PKR-positive dRIFs (Fig. 7F). This is consistent with a recent study that showed that phospho-PKR is only transiently present in dRIFs as PKR rapidly dissociates from dsRNA after phosphorylation [16]. The percentage of cells containing phospho-PKR foci decreased by at least twofold in the presence of most N proteins, except HCoV-OC43 N which had no influence on phospho-PKR foci formation and most often did not colocalize with dsRNA foci (Fig. 7E, F). These data support a model wherein most HCoV N proteins inhibit PKR phosphorylation. In instances where HCoV-OC43 N colocalized with dsRNA, these foci also contained phospho-PKR (Fig. 7E), suggesting that OC43 N is unable to inhibit the phosphorylation of PKR within dRIFs.

### HCoV N proteins alter G3BP1 foci abundance, morphology and composition

In addition to dRIFs, two types of cytosolic biomolecular condensates that form in response to dsRNA include stress granules (SGs) and RNase L-induced bodies (RLBs) [5,6,14]. While SGs and RLBs differ in their morphology and composition, both contain G3BP1 and are induced by dsRNA; thus, we will collectively refer to these as “G3BP1 foci.” SARS-CoV-2 N localizes to G3BP1 foci via a ϕxFG motif within its N-IDR that enables it to directly interact with G3BP1, promoting viral replication [50,55,73,75,76]. The ϕxFG motif is conserved in SARS-CoV-1 N, which also localizes to G3BP1 foci [74], but is not conserved in other HCoV N proteins (Fig. 8A). Thus, we wondered whether HCoV N proteins differ in their modulation of G3BP1 foci.

**Figure 8.**
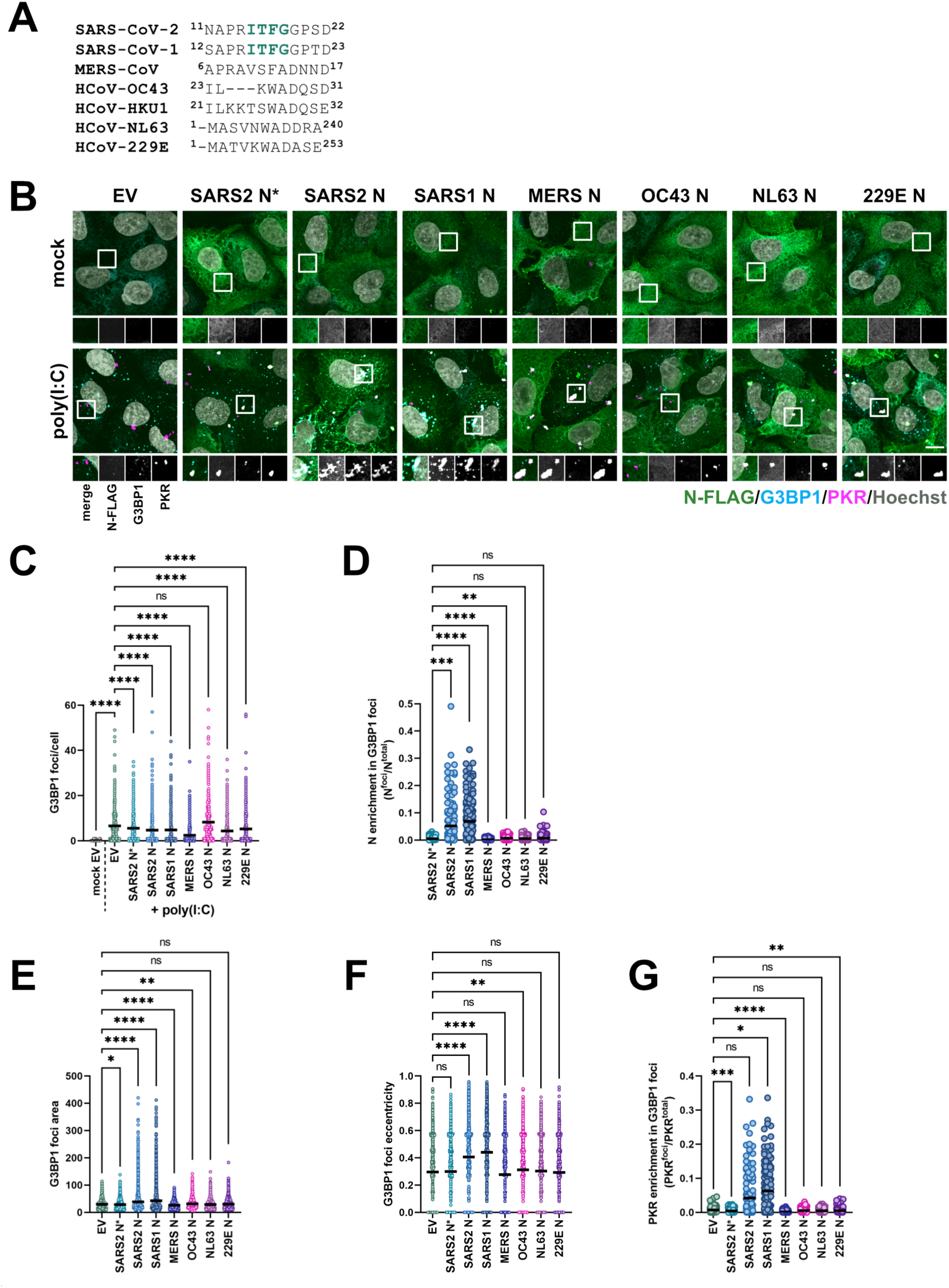
HCoV N proteins differentially alter the morphology and composition of G3BP1 foci. **A.** Sequence conservation of the SARS-CoV-2 G3BP1-binding motif (teal) in HCoV N proteins. **B.** A549 cells were transduced with recombinant lentiviruses expressing an empty vector control (EV) or each of the indicated HCoV N-FLAG constructs and selected with puromycin. Six days post-transduction, cells were mock-transfected or transfected with 0.5 µg poly(I:C). Cells were fixed three hours post-poly(I:C) addition and immunostained with antibodies specific to FLAG (N protein; green), G3BP1 (cyan), or PKR (magenta). Nuclei were stained with Hoechst (grey). Images were obtained using an LSM 880 Airyscan confocal microscope, with maximum-intensity projections presented (scale bar = 10 µm). **C.** Coverslips from B. were imaged using a Zeiss AxioObserver Z1 microscope. The number of G3BP1 foci per cell (for cells transduced with EV) or N-expressing cell (for cells transduced with HCoV N constructs) was quantified in CellProfiler. At least 181 cells were imaged per biological replicate (*n* = 3; mean). Statistics were performed using a Kruskal-Wallis H test with Dunn’s post-hoc analysis (****, p < 0.0001; ns, nonsignificant). **D.** The ratio of N intensity within G3BP1 foci (N^foci^) relative to total N intensity (N^total^) per N-expressing cell in confocal images from B. was quantified in CellProfiler. At least 39 cells were imaged per biological replicate (*n* = 3; mean). Statistics were performed using a Kruskal-Wallis H test with Dunn’s post-hoc analysis (****, p < 0.0001; ***, p < 0.001; **, p < 0.01; ns, nonsignificant). **E-F.** The area (E) and eccentricity (non-roundness) (F) of G3BP1 foci were quantified in CellProfiler using images from B. Each data point represents a single G3BP1 punctum in cells (for cells transduced with EV) or N-expressing cells (for cells transduced with HCoV N constructs). At least 516 puncta were imaged per biological replicate (*n* = 3; mean). Statistics were performed using a Kruskal-Wallis H test with Dunn’s post-hoc analysis (****, p < 0.0001; **, p < 0.01; *, p < 0.05; ns, nonsignificant). **G.** The ratio of PKR intensity within G3BP1 foci (PKR^foci^) relative to total PKR intensity (N^total^) per cell (for cells transduced with EV) or N-expressing cell (for cells transduced with HCoV N constructs) in confocal images from B. was quantified in CellProfiler. At least 39 cells were imaged per biological replicate (*n* = 3; mean). Statistics were performed using a Kruskal-Wallis H test with Dunn’s post-hoc analysis (****, p < 0.0001; ***, p < 0.001; **, p < 0.01; *, p < 0.05; ns, nonsignificant).

To interrogate this, we overexpressed HCoV N proteins in A549 cells, transfected the cells with poly(I:C), and then immunostained for N, G3BP1, and PKR. In the absence of poly(I:C), neither G3BP1 nor PKR formed puncta (Fig. 8B-C). In empty vector-transduced cells, poly(I:C) transfection triggered the formation of both G3BP1 foci and PKR-containing dRIFs. In these control cells, we observed no colocalization between dRIFs and G3BP1 foci, consistent with previous observations made in A549 cells [18] (Fig. 8B). Previous reports suggest that RLBs are the dominant G3BP1 condensate that forms in response to poly(I:C) in A549 cells [14]; consistent with this, the G3BP1 foci we observed in control cells were small and spherical, rather than large and irregular (Fig. 8B). Expression of most HCoV N proteins led to a decrease in the number of G3BP1 foci per cell, with HCoV-OC43 N being the only N protein that had no effect on G3BP1 foci abundance (Fig. 8B-C). This result suggests that HCoV N proteins that inhibit PKR and interact with dsRNA are also able to interfere with the dsRNA-induced formation of G3BP1 foci.

We wondered whether HCoV N proteins interact with G3BP1 foci and/or affect G3BP1 foci composition. As expected, we observed that SARS-CoV-2 and SARS-CoV-1 N, which contain the ϕxFG motif, colocalized with G3BP1 foci, while the other N proteins were unable to do so (Fig. 8B, D). The size and morphology of G3BP1 foci appeared to be altered by the presence of SARS-CoV-2 or SARS-CoV-1 N, as sarbecovirus N-positive G3BP1 foci appeared larger and less circular (Fig. 8E-F). Meanwhile, MERS-CoV, HCoV-NL63, and HCoV-229E N had no effect on the morphology of G3BP1 foci, despite attenuating their formation. Strikingly, while PKR was not present in G3BP1 foci in control cells or in cells expressing MERS-CoV, HCoV-OC43, HCoV-NL63, or HCoV-229E N, PKR was enriched within sarbecovirus N-positive G3BP1 foci (Fig. 8B, G). Together, these findings demonstrate that HCoV N proteins have differential effects on G3BP1 foci. N proteins other than HCoV-OC43 N inhibit G3BP1 foci formation, while sarbecovirus N proteins uniquely alter these foci to relocalize PKR to G3BP1 foci (Fig. 8G).

## DISCUSSION

Previous studies have identified the SARS-CoV-2 N protein as a key player in innate immune antagonism; however, the immune-evasive functions of other HCoV N proteins remain largely unknown. Taken together, these data reveal an unexpected level of heterogeneity in the ability of HCoV N proteins to evade dsRNA responses (Fig. 9). We report five novel findings about immune evasive capabilities of HCoV N proteins. 1) Only highly pathogenic HCoV N proteins can inhibit IFN and OAS/RNase L. 2) Most of the HCoV N proteins tested inhibit PKR, except HCoV-OC43 N. 3) Multiple regions of N, including but not limited to the dsRNA-binding motif and CTD, determine innate immune evasion ability. 4) The ability of N proteins to inhibit PKR correlates with their ability to colocalize and interact with dsRNA, a feature that facilitates colocalization with and reduced activation of PKR. 5) While all HCoV N proteins except HCoV-OC43 N attenuate G3BP1 foci formation, sarbecovirus N proteins are uniquely able to colocalize with G3BP1 foci and relocalize PKR to these foci. The unique dsRNA antagonism abilities of different HCoV N proteins have important implications for our understanding of HCoV replication and pathogenesis, and for the use of common cold HCoVs as BSL-2 proxies for highly pathogenic HCoVs.

**Figure 9.**
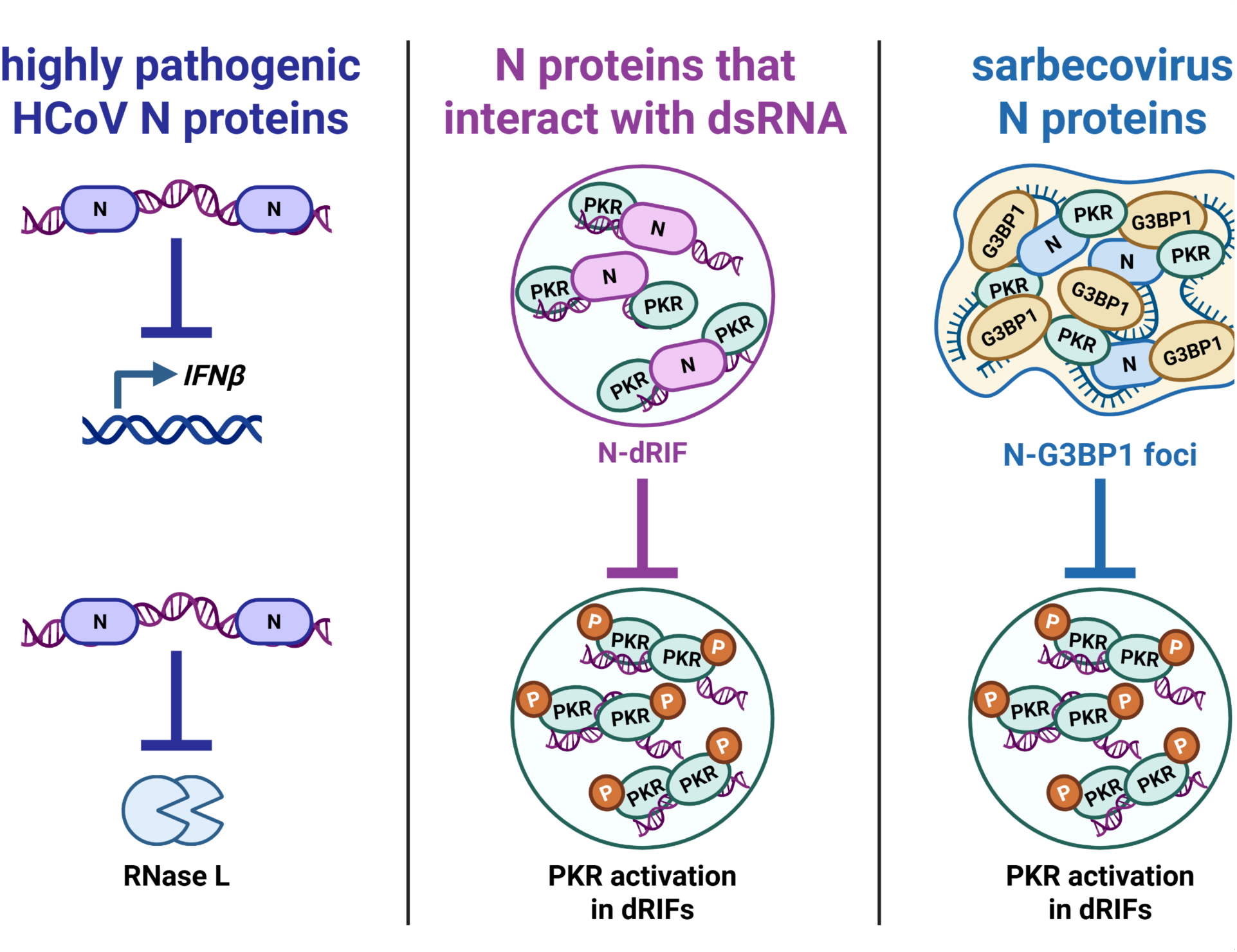
HCoV N proteins differentially antagonize innate immune responses induced by dsRNA. HCoV N proteins antagonize dsRNA responses via multiple mechanisms. Highly pathogenic HCoV N proteins inhibit *IFNβ* induction and RNase L activation in a manner that is likely dependent on their ability to bind dsRNA (left panel). N proteins that interact with dsRNA colocalize with dsRNA and PKR within dRIFs, preventing PKR phosphorylation (middle panel). Sarbecovirus N proteins relocalize PKR to G3BP1 foci, contributing to a decrease in PKR phosphorylation (right panel).

### HCoV N proteins differentially contribute to viral innate immune evasion

HCoVs use multiple strategies to evade dsRNA responses, some universal among HCoVs. For instance, all HCoVs carry out viral RNA synthesis within ER-derived double-membrane vesicles (DMVs) which shield dsRNA intermediates from innate immune sensors [2]. All HCoVs also produce nsp15, an endoribonuclease that limits dsRNA accumulation during infection, preventing activation of IFN, OAS/RNase L, and PKR [27,28,36,77–79]. In addition, some HCoVs produce additional accessory proteins with dedicated innate immune evasive functions. For example, SARS-CoV-2 and SARS-CoV-1 encode ORF6, which inhibits the nucleocytoplasmic trafficking of transcription factors involved in the IFN response [30,32,33].

MERS-CoV produces NS4a, a dsRNA-binding protein that sequesters dsRNA from RLRs, OAS/RNase L and PKR [16,28,29,34,37]. Meanwhile, MERS-CoV and HCoV-OC43 produce NS4b and NS2a, respectively, which are phosphodiesterases that cleave 2-5A, preventing RNase L activation [28,80,81]. We show that the N protein is an important contributor to innate immune evasion for most HCoVs; however, the nature of this contribution differs depending on the virus.

We observed that HCoV-OC43 N was the sole N protein unable to antagonize PKR (Fig. 1G-H); correspondingly, HCoV-229E actively inhibited PKR during infection, while HCoV-OC43 did not (Fig. 2G-J, Fig. 3H-K). This raises the possibility that N proteins may act as determinants of PKR activity during coronavirus infection.

We also observed that highly pathogenic HCoV N proteins inhibited IFN and OAS/RNase L, while common cold HCoV N proteins did not (Fig. 1D-F), yet infection with HCoV-OC43 and HCoV-229E viruses actively inhibited *IFNβ* induction and RNase L activation (Fig 2A-F, Fig. 3B-G). This indicates that these viruses encode additional factors to restrict these pathways. HCoV-OC43 evades RNase L activation using NS2a [80], but it is unclear clear how HCoV-229E inhibits RNase L, and how both viruses inhibit IFN induction. The activity of nsp15 alone is unlikely to explain the blockade of innate immune pathways we observed when infected cells were transfected with poly(I:C), because nsp15 specifically cleaves RNA at uridine residues, which are absent in poly(I:C) [82–84]. Hence, our study suggests that additional common cold coronavirus proteins may have uncharacterized roles in inhibiting IFN and OAS/RNase L.

While it is poorly understood why some HCoVs cause more severe disease than others, one factor thought to influence pathogenesis is the ability of highly pathogenic HCoVs to antagonize IFN responses to a greater extent than common cold HCoVs [85,86]. This difference in IFN antagonism has been previously attributed to the unique accessory proteins encoded by highly pathogenic HCoVs, but our findings raise the intriguing possibility that the N protein may also contribute to differences in HCoV innate immune evasion and pathogenesis. Though we observed that both HCoV-OC43 and HCoV-229E inhibited *IFNβ* induction and OAS/RNase L activation regardless of the activity of their N proteins in cell culture (Fig 2A-F, Fig. 3B-G), it is possible that N could play an important role in innate immune evasion in an organismal context, contributing to disparate HCoV virulence.

### Highly pathogenic HCoV N proteins inhibit IFN and OAS/RNase L independently of their ability to colocalize with dsRNA foci

Previous studies of SARS-CoV-2 N have suggested that its innate immune antagonism ability is conferred primarily by the dsRBM, which was thought to bind dsRNA and prevent its detection by innate immune sensors [49,53,58]. Accordingly, we confirmed that an intact dsRBM is required for inhibition of dsRNA responses by SARS-CoV-2 and HCoV-229E N (Fig. 5).

However, our findings suggest that the dsRBM is not sufficient to confer dsRNA binding ability, as HCoV-OC43 N contains the dsRBM but does not colocalize or coprecipitate with dsRNA (Fig. 4). While the specific features of N that enable dsRNA binding remain undefined, we observed that a chimeric HCoV-OC43 N protein containing the SARS-CoV-2 N carboxy terminus was able to colocalize with dsRNA foci, suggesting that the CTD and C-IDR are major determinants of dsRNA interaction ability (Fig. 6B-C).

Our observations also suggest that colocalization of N with dsRNA is not sufficient to enable IFN and OAS/RNase L antagonism, because HCoV-NL63 and HCoV-229E N were able to interact with dsRNA and/or colocalize with dsRNA foci, but were not able to inhibit IFN and OAS/RNase L (Fig. 1D-F, Fig. 4). Consistent with this, a chimeric HCoV-OC43 N mutant containing the carboxy terminus of HCoV-229E N gained the ability to strongly inhibit IFN and OAS/RNase L, despite poorly colocalizing with dsRNA foci (Fig. 6B-C, Fig. S5). This finding also suggests that while the carboxy terminus of HCoV-229E N may be able to inhibit IFN and OAS/RNase L, the amino terminus of HCoV-229E N could have a dominant negative effect that prevents IFN and OAS/RNase L antagonism.

The mechanism by which highly pathogenic HCoV N proteins inhibit IFN remains enigmatic. Previous studies have suggested that highly pathogenic HCoV N proteins may antagonize IFN by interacting with RIG-I, inhibiting TRIM25-mediated RIG-I ubiquitination, interfering with MAVS activity, and blocking the nuclear translocation of IRF3 and STATs [25,26,31,35,38,39]. While it is unclear why the dsRBM would be required for these processes, it is possible that it plays a role in IFN inhibition independent of its role in dsRNA binding. The mechanism by which highly pathogenic N proteins inhibit OAS/RNase L also remains unresolved. It is possible that these N proteins directly interact with host RNAs including rRNAs, shielding them from cleavage by RNase L. Also, like PKR, OAS and RNase L have been shown to localize to and become activated within dRIFs [19], and it is plausible that highly pathogenic HCoV N proteins interfere with these processes when they localize to dRIFs (Fig. 7). However, if this is the case, it is unclear why HCoV-NL63 and HCoV-229E N proteins, which also localize to dRIFs, do not inhibit RNase L. One possibility is that differences in the biophysical properties and/or phase separation propensity of HCoV N proteins enable only some of them to interfere with OAS/RNase L recruitment to or function within dRIFs [87].

### N proteins that interact with dsRNA impair PKR function within dRIFs

dRIFs are recently discovered biomolecular condensates that contain dsRNA and recruit dsRNA binding proteins including PKR, OAS, PACT, ADAR1, and DHX9 [18–21]. dRIFs are thought to enhance PKR and OAS/RNase L signaling by concentrating these molecules on dsRNA [16–19]; thus, it stands to reason that viruses would target dRIFs in order to prevent activation of these antiviral responses. Recently, Blomqvist et al. (2026) reported that NS4a, a dsRNA binding protein produced by MERS-CoV, condenses on dsRNA when it leaks out of DMVs, competitively inhibiting condensation of PKR on dsRNA and antagonizing PKR activation [16]. Because most HCoV N proteins colocalize with dsRNA, and this ability correlates with their ability to inhibit PKR (Fig. 1G-H, Fig. 4, Fig. 6), we hypothesized that N proteins similarly antagonize PKR by sequestering dsRNA and inhibiting dRIF formation. However, contrary to this hypothesis, we found that N proteins do not block PKR colocalization with dsRNA; instead, they colocalize with PKR and inhibit its activation (Fig. 7). To our knowledge, this is the first time a viral protein has been shown to modulate dRIF function by colocalizing with PKR in dRIFs.

The precise mechanism by which N localization to dRIFs interferes with PKR activation is not clear. One possibility is that N proteins could act as a physical barrier that separates PKR monomers on dsRNA, preventing them from encountering each other and dimerizing. N proteins could also sterically hinder PKR phosphorylation, or prevent PKR from scanning along dsRNA; indeed, the PKR-regulatory protein PACT has recently been shown to inhibit PKR independently of dsRNA sequestration through the latter mechanism [88]. In addition, N proteins could alter PKR activation by affecting dRIF recruitment of other proteins that modulate PKR activity, such as PACT, DHX9, and ZNF346 [20,88]. Finally, N proteins could interact simultaneously with PKR and dsRNA, trapping PKR within dsRNA foci and impeding the typically rapid dissociation of phosphorylated PKR from dsRNA [16]. Supporting this idea, we observed multiple instances where N proteins colocalized with phospho-PKR within dRIFs (Fig. 7E). The consequences of N inhibition of PKR dissociation from dsRNA would be twofold. First, the trapping of phospho-PKR on dsRNA would inhibit eIF2α phosphorylation, because eIF2α is absent from dRIFs and must be phosphorylated in the cytosol [16,21]. Second, immobilization of PKR on a dsRNA would mean that new, non-phosphorylated PKR molecules would be unable to associate with that dsRNA, preventing the amplification of PKR signaling. This would be especially relevant in the context of viral infection, where cytosolic dsRNA is limiting because most dsRNA is sequestered within DMVs. This model is also consistent with our observation that N protein colocalization with dsRNA led to an overall decrease in the abundance of phospho-PKR foci.

### HCoV N proteins differentially modulate G3BP1 foci

In addition to inducing dRIF formation, dsRNA can trigger the formation of other ribonucleoprotein granules containing G3BP1, such as SGs and RLBs [5,6,14,89]. G3BP1 foci are thought to have antiviral activity, in part because they may serve as platforms for antiviral signaling, and because they may be able to condense and translationally repress viral RNA or promote its decay [5,7–13,15]; thus, HCoVs use multiple strategies to inhibit G3BP1 condensation. For instance, the nsp1 proteins of SARS-CoV-2, SARS-CoV-1, and HCoV-OC43, which mediate host shutoff, inhibit SG formation [9]. The full-length N proteins of SARS-CoV-2 and SARS-CoV-1 use the ϕxFG motif within their N-IDR to interact with G3BP1, inhibiting SG/RLB formation and promoting viral replication [9,50–52,54–57,74–76,90], while the amino terminally truncated SARS-CoV-2 N proteoform N* blocks G3BP1 foci formation via a unique mechanism that is independent of PKR activation or G3BP1 binding but requires dsRNA binding [53]. dsRNA binding by NS4a in MERS-CoV and N in SARS-CoV-2 have also been previously shown to inhibit dsRNA-induced G3BP1 foci formation by inhibiting upstream activation of PKR and/or RNase L [34,49,53]. Here, we show that all N proteins except HCoV-OC43 N can inhibit dsRNA-induced formation of G3BP1 foci (Fig. 8C). Because the ability to limit the abundance of G3BP1 foci correlated with the ability of N proteins to inhibit PKR (Fig. 1G-H), we speculate that N predominantly inhibits G3BP1 foci formation in our system via upstream antagonism of PKR.

We observed that sarbecovirus N proteins were the only HCoV N proteins able to colocalize with G3BP1 foci (Fig. 8D), and that this colocalization event altered the morphology of G3BP1 foci to form larger and less spherical granules than canonical RLBs (Fig. 8E-F). The biological implications of these findings are unclear. It is possible that localization of N to G3BP1 foci protects G3BP1-localized RNA from degradation mediated by RNase L or other factors such as nsp1 during infection, consistent with a previous report showing that the G3BP1-N interaction plays a role in enhancing the local translation of viral mRNAs early in SARS-CoV-2 infection [75]. Sarbecovirus N proteins also relocalized PKR from dsRNA-containing dRIFs to G3BP1 foci that lack dsRNA (Fig. 8G). We predict that PKR within N-containing G3BP1 foci is inactive, as it is sequestered away from dsRNA, making this another way for sarbecovirus N proteins to antagonize PKR activation. By explaining how full-length SARS-CoV-2 N can inhibit PKR even without localizing to dRIFs, this finding reconciles the prior confusing observation that full-length SARS-CoV-2 N inhibited PKR to the same extent as the truncated SARS-CoV-2 N* protein, despite the latter having enhanced dsRNA binding ability [53]. We posit that interaction with G3BP1 foci and inhibition of dRIF-localized dsRNA responses represent competing functions of N in sarbecoviruses, because these foci are spatially separated in the cytosol and that sarbecoviruses have evolved two distinct solutions to this trade-off: i) SARS-CoV-2 variants of concern evolved the ability to produce large amounts of N*, which localizes to dRIFs but not G3BP1 foci [53,62], and ii) sarbecovirus N proteins have the ability to divert PKR from dRIFs to G3BP1 foci, allowing them to inhibit PKR and interact with G3BP1 simultaneously.

### Significance

Here, we provide a comprehensive comparison of the abilities of HCoV N proteins to evade the IFN, OAS/RNase L, and PKR pathways. Our findings corroborate existing research which situates the N protein as an essential antagonist of innate immune responses, while also revealing novel key disparities in how the N proteins of different HCoVs block these pathways. These differences in N function may have important consequences for viral replication and pathogenesis; for instance, the enhanced ability of highly pathogenic HCoV N proteins to inhibit IFN and OAS/RNase L may contribute to the more severe disease caused by these viruses. In addition, the heterogeneity we observed between HCoV N proteins suggests that the field should exercise caution when utilizing common cold HCoVs as BSL-2 model systems for studying the replication or antiviral susceptibility of BSL-3 HCoVs. Even in the case of essential structural proteins like N, accessory functions of these proteins are not always conserved. Finally, we reveal novel mechanisms by which HCoV N proteins alter dRIFs and G3BP1 foci, contributing to a growing body of work showing that modulation of host biomolecular condensates is a universal strategy used by viruses to evade antiviral responses and promote their own replication.

### Limitations

Our study has the following limitations. First, we characterized the innate immune evasion abilities of HCoV N proteins using a reductionist N overexpression system. This system does not fully recapitulate the environment of a viral infection, where factors such as the presence of other viral proteins, the posttranslational modification of N by host and viral proteins [51,74,91–99], the induction of host stress and inflammatory responses, and differences in optimal viral infection temperatures [48,85] could affect N function. Second, our overexpression experiments used the synthetic dsRNA poly(I:C), which have properties that differ from the viral dsRNAs that would be produced by HCoVs during infection. Third, our study used N protein sequences corresponding to viral reference strains. Currently circulating variants of SARS-CoV-2 and common cold coronaviruses have been reported to differentially activate dsRNA responses compared to ancestral strains, and additional work is required to determine whether the N proteins of these contemporary viruses differ in their innate immune evasion abilities [100,101]. Fourth, our characterization of N protein function was limited to coronaviruses that infect humans. A previous study showed that the MHV N protein antagonizes OAS/RNase L, but not PKR, suggesting that other embecovirus N proteins may also lack PKR inhibition ability [102]. In addition, a recent study showed that the G3BP1-binding motif is conserved across the N proteins of bat sarbecoviruses and also evolved independently in a clade of bat merbecovirus, raising the possibility that these bat N proteins may also relocalize PKR to G3BP1 foci [103]. Overall, additional work is needed to determine how the N proteins of other animal coronaviruses evade dsRNA responses.

## MATERIALS AND METHODS

### Cell culture

All cells were grown at 5% CO_2_ and 20% O_2_ in 37°C humidified incubators. EA.Hy926 (ATCC, a generous gift from Dr. Robert Rose), A549 (ATCC), Vero E6 (ATCC), and HEK293T (ATCC) cells were cultured in Dulbecco’s Modified Eagle Medium (DMEM; Thermo Fisher) supplemented with 10% heat-inactivated fetal bovine serum (FBS; Thermo Fisher) and penicillin (100 U/mL), streptomycin (100 μg/mL), L-glutamine (2 mM) (PSQ; Thermo Fisher). MRC-5 cells (ATCC, a generous gift from Dr. David Proud) were cultured in Eagle’s Minimum Essential Medium (EMEM) supplemented with 10% FBS (Thermo Fisher) and penicillin (100 U/mL), streptomycin (100 μg/mL), L-glutamine (2 mM) (Thermo Fisher).

### Virus propagation and infection

HCoV-OC43 (ATCC VR-1558) and HCoV-229E (ATCC VR-740) were propagated in Vero E6 and MRC-5 cells, respectively. To propagate viruses, cells were infected at an MOI of 0.01 in serum-free media at 33°C for 1 hour. After adsorption, serum-free media was replaced with media supplemented with 2% FBS and penicillin (100 U/mL), streptomycin (100 μg/mL), L-glutamine (2 mM). Virus-containing supernatants were harvested 4 days post infection for HCoV-OC43 or 3-4 days post-infection for HCoV-229E. Following clearing of the supernatant by centrifugation, viral stocks were aliquoted and stored at −80 °C. Infectious titer of HCoV-OC43 stocks was determined by plaque assay in Vero E6 cells using equal parts 2.4% w/v semi-solid colloidal cellulose overlay (Sigma; prepared in ddH_2_O) and 2× DMEM (Wisent) and penicillin (100 U/mL), streptomycin (100 μg/mL), L-glutamine (2 mM) (PSQ; Thermo Fisher). Cells were fixed 6 days post infection with 4% formaldehyde, then stained with 1% w/v crystal violet (Thermo Fisher). Infectious titers of HCoV-229E stocks, as well as titers of HCoV-OC43 and HCoV-229E from kinetics experiments, were determined by TCID_50_ in MRC-5 cells using the Reed-Muench method, as described previously [104].

For experiments, cells were seeded to achieve a confluence of approximately 80%. Cells were infected using viral inoculum at the indicated multiplicity of infection (MOI; determined in MRC-5 cells) diluted in serum-free media at 33 °C for 1 hour. Following adsorption, serum-free media was replaced with media supplemented with 10% FBS and penicillin (100 U/mL), streptomycin (100 μg/mL), L-glutamine (2 mM).

### Plasmids and cloning

Codon-optimized N genes were subcloned and FLAG-tagged using BamHI and EcoRI restriction sites into pLVX-IRES-puro, a generous gift from the Krogan lab [105]. pLVX-IRES-eF1alpha-SARS-CoV-2-N-FLAG-puro originated from pLVX-EF1alpha-SARS-CoV-2-N-2xStrep-IRES-puro, a generous gift from the Krogan lab [105]. pLVX-IRES-eF1alpha-SARS-CoV-1-N-FLAG-puro originated from VG40143-G-N (Sino Biological). pLVX-IRES-eF1alpha-MERS-CoV-N-FLAG-puro originated from VG40068 (Sino Biological). pLVX-IRES-eF1alpha-HCoV-OC43-N-FLAG-puro, pLVX-IRES-eF1alpha-HCoV-NL63-N-FLAG-puro, and pLVX-IRES-eF1alpha-HCoV-229E-N-FLAG-puro respectively originated from pGBW-m4134906, pGBW-m4134910, or pGBW-m4134902, gifts from Ginkgo Bioworks & Benjie Chen. SARS-CoV-2-N-FL-M210I-M234V-FLAG (referred to as SARS-CoV-2 N within text), SARS-CoV-2-N*^M210^-M234V-FLAG (referred to as SARS-CoV-2 N* within text), and SARS-CoV-2-N-FL-M210I-M234V-K257A-K261A-FLAG (referred to as SARS2 N^ΔRBM^ within text) [53] were subcloned using BamHI and EcoRI restriction sites into pLVX-IRES-puro. HCoV-229E-N-K246R-K250R-FLAG (referred to as 229E N^ΔRBM^ within text) was generated by first subcloning HCoV-229E N into pcDNA3.1(+), then performing site-directed mutagenesis and subcloning the mutant construct using BamHI and EcoRI restriction sites into pLVX-IRES-puro.

HCoV-OC43-N^1-258^-SARS-CoV-2-N^251-419^-FLAG and HCoV-OC43-N^1-258^-HCoV-229E-N^236-389^-FLAG (referred to as OC-SARS2 and OC-229E respectively within text) were generated by overlap extension PCR. The HCoV-OC43 N^1-258^, SARS-CoV-2 N^251-419^-FLAG, and HCoV-229E-N^236-389^-FLAG fragments were amplified with appropriate overlapping overhangs, then HCoV-OC43 N^1-258^ was combined with SARS-CoV-2 N^251-419^-FLAG or HCoV-229E-N^236-389^-FLAG without primers for a second PCR step to create chimeric fragments. Chimeric fragments were amplified using N- and C-terminal flanking primers, subcloned into pcDNA 3.1(+) using BamHI and EcoRI restriction sites, and finally subcloned into pLVX-IRES-puro using BamHI and EcoRI. Plasmids can be found in Table 2, while mutagenesis primers can be found in Table 3.

**Table 2.** Plasmids.

| Plasmid | Use | Source |
| --- | --- | --- |
| pLVX-IRES-puro | Empty vector control | pLVX-IRES-puro was a generous gift from the Krogan lab [105] |
| pLVX-IRES-eF1alpha-SARS-CoV-2-N-FL-M210I-M234V-FLAG-puro | Overexpression | [53] |
| pLVX-IRES-eF1alpha-SARS-CoV-2-N*M210-M234V-FLAG-puro | Overexpression | [53] |
| pLVX-IRES-eF1alpha-SARS-CoV-1-N-FLAG-puro | Overexpression | VG40143-G-N (Sino Biological) |
| pLVX-IRES-eF1alpha-MERS-CoV-N-FLAG-puro | Overexpression | VG40068 (Sino Biological) |
| pLVX-IRES-eF1alpha-HCoV-OC43-N-FLAG-puro | Overexpression | pGBW-m4134906 (Addgene #151960) |
| pLVX-IRES-eF1alpha-HCoV-NL63-N-FLAG-puro | Overexpression | pGBW-m4134910 (Addgene #151939) |
| pLVX-IRES-eF1alpha-HCoV-229E-N-FLAG-puro | Overexpression | pGBW-m4134902 (Addgene #151912) |
| pLVX-IRES-eF1alpha-SARS-CoV-2-N-FL-M210I-M234V-K257A-K261A-FLAG-puro | Overexpression | [53] |
| pLVX-IRES-eF1alpha-HCoV-229E-N-K246R-K250R-FLAG-puro | Overexpression | HCoV-229E-N-FLAG |
| pLVX-IRES-eF1alpha-HCoV-OC43-N<sup>1-258</sup>-SARS-CoV-2-N<sup>251-419</sup>-FLAG | Overexpression | HCoV-OC43-N-FLAG, SARS-CoV-2-N-FL-FLAG |
| pLVX-IRES-eF1alpha-HCoV-OC43-N<sup>1-258</sup>-HCoV-229E-N<sup>236-389</sup>-FLAG | Overexpression | HCoV-OC43-N-FLAG, HCoV-229E-N-FLAG |
| pMD2.G | Lentivirus generation | Addgene #12259 |
| psPAX2 | Lentivirus generation | Addgene #12260 |

**Table 3.**
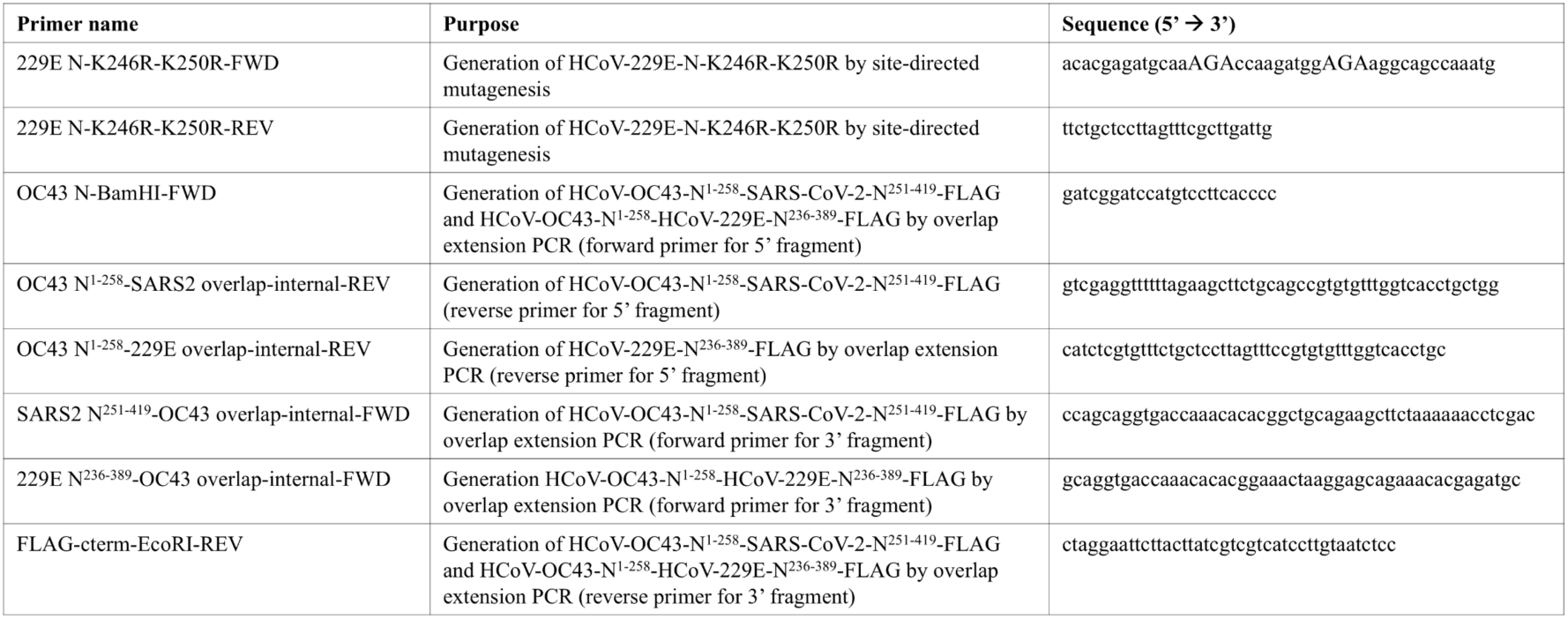
Mutagenesis primers.

### Production and use of recombinant lentiviruses

Second-generation recombinant lentiviruses were generated as in [104]. Briefly, HEK293T cells were seeded on a 10-cm dish to achieve a confluence of approximately 80%. pSPAX2 and pMD2.G (gifts from Didier Trono), and the appropriate lentiviral transfer plasmid were combined in Opti-MEM (Gibco) at a 2:1:3.3 µg ratio, then mixed with linear polyethylenimine (1 mg/mL PEI MAX, Polysciences) in Opti-MEM at a 1 µg DNA: 3 µL PEI ratio. Transfection mixture was added dropwise to cells containing serum-free media, then replaced with antibiotic-free media containing 10% FBS, 2 mM L-glutamine six hours post-transfection. Virus-containing supernatant was harvested and filtered with a 0.45 µm syringe filter (VWR) 48 hours post-transfection. Filtered supernatant was aliquoted and stored at −80 °C for single use. For each N-expressing lentivirus stock, appropriate lentivirus dilutions to achieve approximately equal levels of protein expression were determined by transduction followed by immunoblotting.

For transductions, cells were seeded on 6-well plates to achieve a confluence of approximately 40%. Lentiviruses were thawed at 37 °C, then diluted in complete media containing 5 µg/mL polybrene (Sigma). 24 hours post-transduction, cells were expanded into 10-cm dishes, and the following day, media was replaced with complete media containing 1 µg/mL puromycin (Thermo Fisher). Puromycin-containing media was replaced with complete media 24-30 hours post-transduction. Following 24 hours of recovery, cells were seeded onto 12-well plates for experiments. For overexpression of SARS-CoV-2 N and SARS-CoV-2 N*, constructs were used in which internal methionine residues were mutated to ensure that only the full-length N proteoform was expressed (M210I for SARS-CoV-2 N*; M210I and M234V for full-length SARS-CoV-2 N) as in [53].

### Poly(I:C) transfections

Poly(I:C) transfections were carried out using FuGene HD (Promega) according to manufacturer’s instructions. Briefly, 0.5 or 1µg high molecular weight poly(I:C) (for a 12-well or 6-well plate, respectively) were combined with FuGene HD at a 1 µg poly(I:C): 3 µL FuGene ratio in Opti-MEM (Gibco), and the transfection mixture was added dropwise to cells growing in complete media.

### Reverse transcriptase quantitative PCR

RNA was harvested with the RNeasy Mini kit (Qiagen) according to manufacturer’s instructions and stored at −80 °C. To generate cDNA, RNA concentrations were determined by NanoDrop One^C^ (Thermo Fisher), then 500 ng RNA was converted to cDNA with Maxima H Minus Reverse Transcriptase (Thermo Fisher) according to manufacturer’s instructions. RT-qPCR was carried out as described in [53]. cDNA was diluted 1:10 in nuclease-free water and amplification was performed using SsoFast EvaGreen Master Mix (Bio-Rad). RT-qPCR cycling conditions: *initial denaturation at 98 °C - 2:00 min; cycling at 39 × (98 °C - 0.02 min; annealing and amplification at 60 °C - 0.05 min); final extension at 65 °C - 0.10 min, followed by melt curve analysis (60 °C - 0.10 min and 95 °C - 0.2 min).* Relative RNA abundance was calculated using the 2^−ΔΔCt^ equation, with 18S rRNA being used as a housekeeping gene for normalization. qPCR primers are listed in Table 4.

**Table 4.**
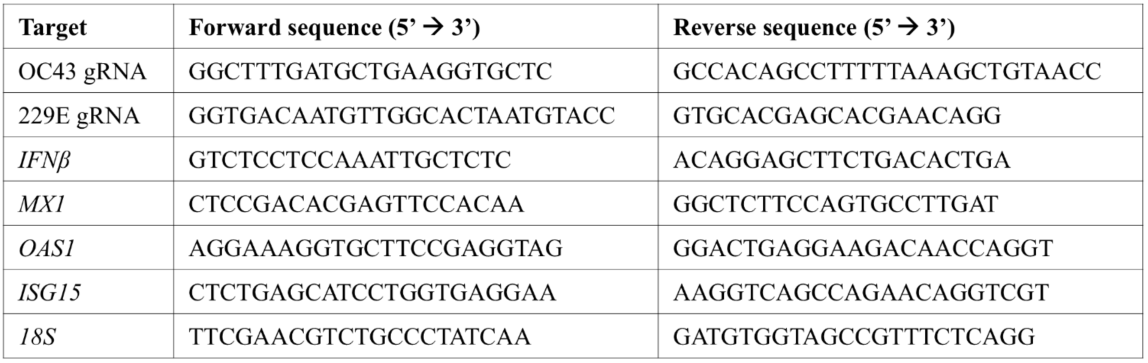
qPCR primers.

### rRNA integrity assay

Cells were transfected with 0.5 µg high molecular weight poly I:C (Invitrogen) for 3 hours, and intracellular RNA was harvested using RNeasy Mini kit (Qiagen) according to manufacturer’s instructions, then stored at −80 °C until use. RNA concentrations were determined using NanoDrop One^C^ (Thermo Fisher) and extracted RNA was diluted to a final concentration of 50 ng/µL, then submitted to the University of Calgary Centre for Health Genomics and Informatics for automated electrophoresis using an Agilent 4200 TapeStation system.

### Immunoblotting and densitometric analysis

Cells were lysed in 1x Laemmli buffer (31.5 mM Tris-HCl pH 6.8, 10% glycerol, 1% SDS), and protein lysates were stored at −20 °C until use. Protein concentration was quantified using DC Protein Assay kit (Bio-Rad) according to manufacturer’s instructions and protein concentrations were normalized by diluting in 1x Laemmli buffer. Protein lysates were boiled at 95 °C for five minutes in the presence of 100 mM DTT (Sigma-Aldrich), then resolved by SDS-PAGE with TGX Stain-Free acrylamide gels, which contain a trihalo compound that enables total protein visualization (Bio-Rad). Proteins were transferred onto PVDF membranes using a Trans-Blot Turbo transfer apparatus (Bio-Rad), and total protein images were acquired using the ChemiDoc MP Imaging system (Bio-Rad). PVDF membranes were blocked with 5% BSA (Thermo-Fisher) diluted in TBST (10 mM Tris-HCl pH 7.4, 150 mM NaCl, 0.1% Tween-20). Primary antibodies were diluted in 5% BSA in TBST, and membranes were incubated with primary antibodies overnight at 4 °C. Secondary antibodies were diluted in 5% skim milk in TBST, except for anti-phospho-PKR and anti-phospho-eIF2α antibodies, which were diluted in 5% BSA in TBST. Membranes were incubated with secondary antibodies for one hour at room temperature, then images were acquired using the ChemiDoc MP Imaging system (Bio-Rad). Antibodies used for immunoblotting are listed in Table 5.

**Table 5.** Antibodies.

| Antibody | Species | Vendor/Catalog # | Application | Dilution |
| --- | --- | --- | --- | --- |
| FLAG | Rabbit | Cell Signaling Technology/14793S | Immunoblotting<br>Immunofluorescence | 1:1000<br>1:1000 |
| FLAG | Mouse | Cell Signaling Technology/8146S | Immunoprecipitation | 1:100 |
| FLAG | Rat | Novus Biologicals/NBP1-06712 | Immunofluorescence | 1:100 |
| PKR | Rabbit | Cell Signaling Technology/12297S | Immunoblotting<br>Immunofluorescence | 1:1000<br>1:500 |
| p-PKR | Rabbit | Abcam/ab32036 | Immunoblotting | 1:1000 |
| p-PKR | Rabbit | Thermo Fisher/MA5-32086 | Immunofluorescence | 1:500 |
| eIF2α | Rabbit | Cell Signaling Technology/9722S | Immunoblotting | 1:1000 |
| p-eIF2α | Rabbit | Cell Signaling Technology//9721S | Immunoblotting | 1:1000 |
| HCoV-OC43 N | Mouse | Sigma-Aldrich/MAB9012 | Immunoblotting | 1:1000 |
| HCoV-229E N | Rabbit | Sino Biological/40640-T62 | Immunoblotting | 1:5000 |
| K1 (dsRNA) | Mouse | Cell Signaling Tehcnology/28764S | Immunofluorescence | 1:1000 |
| G3BP1 | Mouse | BD Bioscences/611126 | Immunofluorescence | 1:500 |
| Alexa Fluor 488 anti-rabbit IgG | Chicken | Thermo Fisher/A-21441 | Immunofluorescence | 1:1000 |
| Alexa Fluor 488 anti-rat IgG | Goat | Thermo Fisher/A-1106 | Immunofluorescence | 1:500 |
| Alexa Fluor 555 anti-mouse IgG | Donkey | Thermo Fisher/A-31570 | Immunofluorescence | 1:1000 |
| Alexa Fluor 647 anti-mouse IgG | Chicken | Thermo Fisher/A-21463 | Immunofluorescence | 1:1000 |
| Alexa Fluor 647 anti-rabbit IgG | Chicken | Thermo Fisher/A-21443 | Immunofluorescence | 1:1000 |
| HRP anti-mouse IgG | Horse | Cell Signaling Technology/7076S | Immunoblotting | 1:3333 |
| HRP anti-rabbit IgG | Goat | Cell Signaling Technology/7074S | Immunoblotting | 1:3333 |

### Poly(I:C)-biotin pulldowns

Poly(I:C)-biotin pulldowns were conducted as in [53]. Briefly, A549 cells were transduced in 6-well plates, then were seeded onto 10-cm dishes, selected, and recovered in complete media. 24 hours post-recovery, the media was removed. Cells were washed with ice-cold 1x PBS, then harvested in ice-cold 1x PBS. Cells were subjected to a second PBS wash, then resuspended in lysis buffer (150 mM NaCl, 10 mM Tris pH 7.4, 1 mM EDTA, 1% v/v Triton X-100, 0.5% v/v NP-40, 0.1% v/v P8340 protease inhibitor cocktail (Sigma Aldrich)). Lysates were treated with 1 U/μL RNase T1 (Thermo Fisher) and rocked for 30 min at room temperature, then rotated at 4 °C for 30 minutes. Cell debris was then removed from lysates by centrifugation. M270 Streptavidin magnetic Dynabeads (Thermo Fisher) were prepared and conjugated to 300 ng poly I:C (HMW)-biotin (Invivogen) as in [53], or left unconjugated as a control. Half of the cell lysate volume was combined with conjugated beads, while the other half was combined with unconjugated beads, and then lysates were rotated overnight at 4 °C. Beads were washed three times with wash buffer (150 mM NaCl, 10 mM Tris pH 7.4, 1 mM EDTA), then samples were eluted by boiling beads in 4× Laemmli buffer (Bio-Rad) with 10% v/v β-mercaptoethanol.

### Co-immunoprecipitations

Co-immunoprecipitations were performed as in [106]. Briefly, A549 cells were transduced in 6-well plates, then selected and recovered in complete media. 24 hours post-recovery, cells were transfected with 1 μg poly(I:C) or mock-transfected for three hours. Cells were washed in ice-cold 1x PBS, then harvested in ice-cold 1x PBS. Following a second PBS wash, cells were resuspended in lysis buffer (150 mM NaCl, 10 mM Tris pH 7.4, 1 mM EDTA, 1% v/v Triton X-100, 0.5% v/v NP-40, Roche protease inhibitor tablet), rotated at 4 °C for 30 minutes, and then subjected to centrifugation to remove cell debris. Lysates were incubated with an anti-FLAG antibody (Table 5) overnight at 4°C. Meanwhile, Protein G magnetic Dynabeads (Thermo Fisher) were incubated in blocking buffer (150 mM NaCl, 10 mM Tris pH 7.4, 1 mM EDTA, 5 mg/mL BSA) overnight at 4 °C. Following overnight incubation, beads and lysates were combined for one hour at 4°C, then washed three times with wash buffer (150 mM NaCl, 10 mM Tris pH 7.4, 1 mM EDTA) and boiled in 4× Laemmli buffer (Bio-Rad) with 10% v/v β-mercaptoethanol.

### Immunofluorescence

Cells were seeded onto 18 mm glass round-bottom #1.5 coverslips (Electron Microscopy Sciences), then fixed at experimental endpoint with 4% paraformaldehyde (Electron Microscopy Sciences) in 1x PBS (Gibco) for 10 minutes at room temperature. Permeabilization was carried out with 0.1% (v/v) Triton X-100 (Sigma-Aldrich) diluted in 1× PBS (Gibco) for 10 min at room temperature, then blocking was carried out with 1% (v/v) human AB serum (Sigma-Aldrich) diluted in 1x PBS for one hour at room temperature. Coverslips were incubated overnight at 4 °C with primary antibodies diluted in 1% human AB blocking buffer, then for one hour at room temperature with secondary antibodies and 1 µg/mL Hoechst (Invitrogen) prior to mounting with ProLong Gold Antifade mountant (Thermo Fisher). For three-colour imaging, cells were co-stained with anti-rabbit Alexa Fluor 488 and anti-mouse Alexa Fluor 647 to minimize bleed-through. For four-colour imaging, cells were co-stained with anti-rat Alexa Fluor 488, anti-mouse Alexa Fluor 555, and anti-rabbit Alexa Fluor 647. Images used for colocalization analysis and representative images were acquired using a Zeiss LSM 880 confocal microscope with a 63× oil-immersion objective. Exposure time and laser power were not changed within replicates. For enumeration of PKR, phospho-PKR, and G3BP1 foci, images were acquired using a Zeiss AxioObserver Z1 microscope with a 40× oil-immersion objective. Antibodies used for immunofluorescence are listed in Table 5.

### Image analysis

G3BP1, PKR, and phospho-PKR foci were quantified using CellProfiler [107] as in [53]. Briefly, nuclei were segmented using the “IdentifyPrimaryObjects” module, and a propagation function was applied to each nucleus to define cell borders. Thresholding was applied to identify N-positive cells. G3BP1, PKR, or phospho-PKR foci were enhanced using the “EnhanceOrSuppressFeatures” module, then segmented using the “IdentifyPrimaryObjects” module. The size range for PKR or phospho-PKR foci was defined as 5-35 pixels; meanwhile, the size range for G3BP1 foci was defined as 2-35 pixels to account for size variability between SGs and RLBs. For G3BP1 foci, the “MeasureObjectSizeShape” module was used to quantify foci area and eccentricity (non-roundness). Foci abundance was represented by graphing the number of foci per cell, while area and eccentricity was represented by graphing per-foci values.

For colocalization analysis, images were acquired using a Zeiss LSM 880 confocal microscope with z-stacks. Nuclei, cells, and N-positive cells were identified as above. Enrichment analysis was performed as in [53]. Briefly, to quantify the enrichment of a given molecule (referred to here as “molecule A”; either N or PKR) within foci containing another molecule (referred to here as “molecule B”; either dsRNA, G3BP1 or PKR), a mask was used to define the area within and outside foci containing molecule B. The mean integrated intensity of molecule A within and outside foci of molecule B was then quantified. Percent enrichment of molecule A within foci containing molecule B was calculated by dividing the mean integrated intensity of molecule A within foci (A^foci^) by the sum of the mean integrated intensities of molecule A within and outside foci (A^total^). Enrichment was represented by graphing per cell. For all imaging experiments, quantification parameters were kept identical within replicates, but thresholds were altered between experiments to account for variability in staining.

### Multiple sequence alignment

HCoV N protein sequences were retrieved from the NCBI nucleotide database. Multiple sequence alignment was performed in Clustal Omega [108,109] using default settings. The multiple sequence alignment was visualized in Jalview [110], and sequences were colour coded based on domain predictions obtained from InterPro [111]. N sequence accession numbers are listed in Table 6.

**Table 6.** N sequence accession numbers.

| Coronavirus | N protein accession number (NCBI) |
| --- | --- |
| SARS-CoV-2 | YP_009724397.2 |
| SARS-CoV-1 | YP_009825061.1 |
| MERS-CoV | YP_009047211.1 |
| HCoV-OC43 | YP_009555245.1 |
| HCoV-HKU1 | YP_173242.1 |
| HCoV-NL63 | YP_003771.1 |
| HCoV-229E | NP_073556.1 |

### Statistics

Statistical analyses were performed using GraphPad Prism 9.0. For per-cell foci counts, per-cell enrichment quantifications, and per-puncta shape measurements, independent biological replicates were combined and plotted on a single graph. These per-cell and per-puncta data were non-parametric; thus, statistics were performed on these data using a Kruskal-Wallis test. All other statistics were performed using ANOVA.

## ACKNOWLEDGEMENTS

We thank Dr. Denys Khaperskyy (Dalhousie University) and Dr. James M. Burke (University of Florida Scripps Institute) for insightful comments about this work and Drs. Anne Vaahtokari and Luc Provencher of the Charbonneau Microscopy Facility for microscopy support. We acknowledge all the members of the Corcoran lab for helpful discussions about experimental design. NS was supported by a Master’s Canada Graduate Scholarship from the Canadian Institutes of Health Research, a Cumming School of Medicine graduate training award, an Alberta Graduate Excellence Scholarship, and a University of Calgary of Calgary Silver Anniversary Graduate Fellowship. RPM was supported by a Snyder Institute Beverley Phillips Doctoral training award and a Doctoral Canada Graduate Scholarship from the Canadian Institutes of Health Research. This study was supported by operating funds awarded to JAC from the Canadian Institutes for Health Research (CIHR) (project grant #195645). Schematics were created using BioRender.

## AUTHOR CONTRIBUTIONS

**Noga Sharlin:** Conceptualization; Data curation; Formal analysis; Investigation; Methodology; Validation; Visualization; Writing – original draft; Writing – review & editing

**Rory P. Mulloy:** Data curation; Investigation; Methodology; Writing – review & editing

**Madeline Day:** Data curation; Investigation; Methodology

**Jennifer A. Corcoran:** Conceptualization; Funding acquisition; Project administration; Resources; Supervision; Writing – original draft; Writing – review & editing

**Competing interests:** The authors have no competing interests to declare.

## Notes

### Competing Interest Statement

The authors have declared no competing interest.

## REFERENCES

1. Chen YG, Hur S. Cellular origins of dsRNA, their recognition and consequences. Nat Rev Mol Cell Biol. 2022;23: 286–301. doi:10.1038/s41580-021-00430-1

2. Romero-Brey I, Bartenschlager R. Membranous Replication Factories Induced by Plus-Strand RNA Viruses. Viruses. 2014;6: 2826–2857. doi:10.3390/v6072826

3. Weber F, Wagner V, Rasmussen SB, Hartmann R, Paludan SR. Double-Stranded RNA Is Produced by Positive-Strand RNA Viruses and DNA Viruses but Not in Detectable Amounts by Negative-Strand RNA Viruses. Journal of Virology. 2006;80: 5059–5064. doi:10.1128/jvi.80.10.5059-5064.2006

4. Tanneti NS, Stillwell HA, Weiss SR. Human coronaviruses: activation and antagonism of innate immune responses. Microbiology and Molecular Biology Reviews. 2024;0: e00016–23. doi:10.1128/mmbr.00016-23

5. McCormick C, Khaperskyy DA. Translation inhibition and stress granules in the antiviral immune response. Nat Rev Immunol. 2017;17: 647–660. doi:10.1038/nri.2017.63

6. Protter DSW, Parker R. Principles and Properties of Stress Granules. Trends in Cell Biology. 2016;26: 668–679. doi:10.1016/j.tcb.2016.05.004

7. Brownsword MJ, Locker N. A little less aggregation a little more replication: Viral manipulation of stress granules. WIREs RNA. 2023;14: e1741. doi:10.1002/wrna.1741

8. Burke JM, Ratnayake OC, Watkins JM, Perera R, Parker R. G3BP1-dependent condensation of translationally inactive viral RNAs antagonizes infection. Science Advances. 2024;10: eadk8152. doi:10.1126/sciadv.adk8152

9. Dolliver SM, Kleer M, Bui-Marinos MP, Ying S, Corcoran JA, Khaperskyy DA. Nsp1 proteins of human coronaviruses HCoV-OC43 and SARS-CoV2 inhibit stress granule formation. PLoS Pathog. 2022;18: e1011041. doi:10.1371/journal.ppat.1011041

10. Liu H, Bai Y, Zhang X, Gao T, Liu Y, Li E, et al. SARS-CoV-2 N Protein Antagonizes Stress Granule Assembly and IFN Production by Interacting with G3BPs to Facilitate Viral Replication. Journal of Virology. 2022;96: e00412–22. doi:10.1128/jvi.00412-22

11. Onomoto K, Jogi M, Yoo J-S, Narita R, Morimoto S, Takemura A, et al. Critical Role of an Antiviral Stress Granule Containing RIG-I and PKR in Viral Detection and Innate Immunity. PLOS ONE. 2012;7: e43031. doi:10.1371/journal.pone.0043031

12. Reineke LC, Kedersha N, Langereis MA, van Kuppeveld FJM, Lloyd RE. Stress Granules Regulate Double-Stranded RNA-Dependent Protein Kinase Activation through a Complex Containing G3BP1 and Caprin1. mBio. 2015;6: 10.1128/mbio.02486-14. doi:10.1128/mbio.02486-14

13. Shang Z, Zhang S, Wang J, Zhou L, Zhang X, Billadeau DD, et al. TRIM25 predominately associates with anti-viral stress granules. Nat Commun. 2024;15: 4127. doi:10.1038/s41467-024-48596-4

14. Burke JM, Lester ET, Tauber D, Parker R. RNase L promotes the formation of unique ribonucleoprotein granules distinct from stress granules. Journal of Biological Chemistry. 2020;295: 1426–1438. doi:10.1074/jbc.RA119.011638

15. Watkins JM, Burke JM. RNase L-induced bodies sequester subgenomic flavivirus RNAs to promote viral RNA decay. Cell Reports. 2024;43: 114694. doi:10.1016/j.celrep.2024.114694

16. Blomqvist EK, Bracci N, Winstone H, Watkins JM, Weiss SR, Burke JM. PKR condensation at viral replication complexes initiates its activation. Cell Reports. 2026;45. doi:10.1016/j.celrep.2026.117290

17. Briggs S, Blomqvist EK, Cuellar A, Correa D, Burke JM. Condensation of human OAS proteins initiates diverse antiviral activities in response to West Nile virus. Genes Dev. 2025 [cited 13 Feb 2026]. doi:10.1101/gad.352725.125

18. Corbet GA, Burke JM, Bublitz GR, Tay JW, Parker R. dsRNA-induced condensation of antiviral proteins modulates PKR activity. Proceedings of the National Academy of Sciences. 2022;119: e2204235119. doi:10.1073/pnas.2204235119

19. Cusic R, Burke JM. Condensation of RNase L promotes its rapid activation in response to viral infection in mammalian cells. Science Signaling. 2024;17: eadi9844. doi:10.1126/scisignal.adi9844

20. Loucas G, Parker R. Analysis of dRIF Composition Identifies ZNF346 as a Regulator of PKR Activation. bioRxiv; 2026. p. 2026.06.03.729416. doi:10.64898/2026.06.03.729416

21. Zappa F, Muniozguren NL, Wilson MZ, Costello MS, Ponce-Rojas JC, Acosta-Alvear D. Signaling by the integrated stress response kinase PKR is fine-tuned by dynamic clustering. Journal of Cell Biology. 2022;221: e202111100. doi:10.1083/jcb.202111100

22. Wang B, Wang Y, Pan T, Zhou L, Ran Y, Zou J, et al. Targeting a key disulfide linkage to regulate RIG-I condensation and cytosolic RNA-sensing. Nat Cell Biol. 2025;27: 817–834. doi:10.1038/s41556-025-01646-5

23. Steiner S, Kratzel A, Barut GT, Lang RM, Aguiar Moreira E, Thomann L, et al. SARS-CoV-2 biology and host interactions. Nat Rev Microbiol. 2024;22: 206–225. doi:10.1038/s41579-023-01003-z

24. V’kovski P, Kratzel A, Steiner S, Stalder H, Thiel V. Coronavirus biology and replication: implications for SARS-CoV-2. Nat Rev Microbiol. 2021;19: 155–170. doi:10.1038/s41579-020-00468-6

25. Chang C-Y, Liu HM, Chang M-F, Chang SC. Middle East Respiratory Syndrome Coronavirus Nucleocapsid Protein Suppresses Type I and Type III Interferon Induction by Targeting RIG-I Signaling. Journal of Virology. 2020;94: 10.1128/jvi.00099-20. doi:10.1128/jvi.00099-20

26. Chen K, Xiao F, Hu D, Ge W, Tian M, Wang W, et al. SARS-CoV-2 Nucleocapsid Protein Interacts with RIG-I and Represses RIG-Mediated IFN-β Production. Viruses. 2021;13: 47. doi:10.3390/v13010047

27. Chi X, Liang X, Vaddadi K, Zhang X, Gandikota C, More S, et al. SARS-CoV-2 Nsp15 endoribonuclease subverts host defenses to enhance viral fitness in lung cells. Journal of Virology. 2025;99: e01175–25. doi:10.1128/jvi.01175-25

28. Comar CE, Otter CJ, Pfannenstiel J, Doerger E, Renner DM, Tan LH, et al. MERS-CoV endoribonuclease and accessory proteins jointly evade host innate immunity during infection of lung and nasal epithelial cells. Proceedings of the National Academy of Sciences. 2022;119: e2123208119. doi:10.1073/pnas.2123208119

29. Comar CE, Goldstein SA, Li Y, Yount B, Baric RS, Weiss SR. Antagonism of dsRNA-Induced Innate Immune Pathways by NS4a and NS4b Accessory Proteins during MERS Coronavirus Infection. mBio. 2019;10: 10.1128/mbio.00319-19. doi:10.1128/mbio.00319-19

30. Frieman M, Yount B, Heise M, Kopecky-Bromberg SA, Palese P, Baric RS. Severe Acute Respiratory Syndrome Coronavirus ORF6 Antagonizes STAT1 Function by Sequestering Nuclear Import Factors on the Rough Endoplasmic Reticulum/Golgi Membrane. Journal of Virology. 2007;81: 9812–9824. doi:10.1128/jvi.01012-07

31. Hu Y, Li W, Gao T, Cui Y, Jin Y, Li P, et al. The Severe Acute Respiratory Syndrome Coronavirus Nucleocapsid Inhibits Type I Interferon Production by Interfering with TRIM25-Mediated RIG-I Ubiquitination. Journal of Virology. 2017;91: e02143–16. doi:10.1128/JVI.02143-16

32. Kehrer T, Cupic A, Ye C, Yildiz S, Bouhaddou M, Crossland NA, et al. Impact of SARS-CoV-2 ORF6 and its variant polymorphisms on host responses and viral pathogenesis. Cell Host & Microbe. 2023;31: 1668–1684.e12. doi:10.1016/j.chom.2023.08.003

33. Miorin L, Kehrer T, Sanchez-Aparicio MT, Zhang K, Cohen P, Patel RS, et al. SARS-CoV-2 Orf6 hijacks Nup98 to block STAT nuclear import and antagonize interferon signaling. Proceedings of the National Academy of Sciences. 2020;117: 28344–28354. doi:10.1073/pnas.2016650117

34. Nakagawa K, Narayanan K, Wada M, Makino S. Inhibition of Stress Granule Formation by Middle East Respiratory Syndrome Coronavirus 4a Accessory Protein Facilitates Viral Translation, Leading to Efficient Virus Replication. Journal of Virology. 2018;92: 10.1128/jvi.00902-18. doi:10.1128/jvi.00902-18

35. Oh SJ, Shin OS. SARS-CoV-2 Nucleocapsid Protein Targets RIG-I-Like Receptor Pathways to Inhibit the Induction of Interferon Response. Cells. 2021;10: 530. doi:10.3390/cells10030530

36. Otter CJ, Bracci N, Parenti NA, Ye C, Asthana A, Blomqvist EK, et al. SARS-CoV-2 nsp15 endoribonuclease antagonizes dsRNA-induced antiviral signaling. Proceedings of the National Academy of Sciences. 2024;121: e2320194121. doi:10.1073/pnas.2320194121

37. Rabouw HH, Langereis MA, Knaap RCM, Dalebout TJ, Canton J, Sola I, et al. Middle East Respiratory Coronavirus Accessory Protein 4a Inhibits PKR-Mediated Antiviral Stress Responses. PLOS Pathogens. 2016;12: e1005982. doi:10.1371/journal.ppat.1005982

38. Wang S, Dai T, Qin Z, Pan T, Chu F, Lou L, et al. Targeting liquid–liquid phase separation of SARS-CoV-2 nucleocapsid protein promotes innate antiviral immunity by elevating MAVS activity. Nat Cell Biol. 2021;23: 718–732. doi:10.1038/s41556-021-00710-0

39. Zhao Y, Sui L, Wu P, Wang W, Wang Z, Yu Y, et al. A dual-role of SARS-CoV-2 nucleocapsid protein in regulating innate immune response. Sig Transduct Target Ther. 2021;6: 1–14. doi:10.1038/s41392-021-00742-w

40. Masters PS. Coronavirus genome packaging and nucleocapsid assembly. Journal of Virology. 2026;0: e01330–25. doi:10.1128/jvi.01330-25

41. McBride R, van Zyl M, Fielding BC. The Coronavirus Nucleocapsid Is a Multifunctional Protein. Viruses. 2014;6: 2991–3018. doi:10.3390/v6082991

42. Wu W, Cheng Y, Zhou H, Sun C, Zhang S. The SARS-CoV-2 nucleocapsid protein: its role in the viral life cycle, structure and functions, and use as a potential target in the development of vaccines and diagnostics. Virol J. 2023;20: 6. doi:10.1186/s12985-023-01968-6

43. Zhang B, Tian J, Zhang Q, Xie Y, Wang K, Qiu S, et al. Comparing the Nucleocapsid Proteins of Human Coronaviruses: Structure, Immunoregulation, Vaccine, and Targeted Drug. Frontiers in Molecular Biosciences. 2022;9. Available: https://www.frontiersin.org/articles/10.3389/fmolb.2022.761173

44. Sola I, Almazán F, Zúñiga S, Enjuanes L. Continuous and Discontinuous RNA Synthesis in Coronaviruses. Annu Rev Virol. 2015;2: 265–288. doi:10.1146/annurev-virology-100114-055218

45. Matsuo T. Viewing SARS-CoV-2 Nucleocapsid Protein in Terms of Molecular Flexibility. Biology. 2021;10: 454. doi:10.3390/biology10060454

46. Peng Y, Du N, Lei Y, Dorje S, Qi J, Luo T, et al. Structures of the SARS-CoV-2 nucleocapsid and their perspectives for drug design. EMBO J. 2020;39: e105938. doi:10.15252/embj.2020105938

47. Grossoehme NE, Li L, Keane SC, Liu P, Dann CE, Leibowitz JL, et al. Coronavirus N Protein N-Terminal Domain (NTD) Specifically Binds the Transcriptional Regulatory Sequence (TRS) and Melts TRS-cTRS RNA Duplexes. Journal of Molecular Biology. 2009;394: 544–557. doi:10.1016/j.jmb.2009.09.040

48. Roden CA, Dai Y, Giannetti CA, Seim I, Lee M, Sealfon R, et al. Double-stranded RNA drives SARS-CoV-2 nucleocapsid protein to undergo phase separation at specific temperatures. Nucleic Acids Research. 2022;50: 8168–8192. doi:10.1093/nar/gkac596

49. Aloise C, Schipper JG, Vliet A van, Oymans J, Donselaar T, Hurdiss DL, et al. SARS-CoV-2 nucleocapsid protein inhibits the PKR-mediated integrated stress response through RNA-binding domain N2b. PLOS Pathogens. 2023;19: e1011582. doi:10.1371/journal.ppat.1011582

50. Biswal M, Lu J, Song J. SARS-CoV-2 Nucleocapsid Protein Targets a Conserved Surface Groove of the NTF2-like Domain of G3BP1. J Mol Biol. 2022;434: 167516. doi:10.1016/j.jmb.2022.167516

51. Cai T, Yu Z, Wang Z, Liang C, Richard S. Arginine methylation of SARS-Cov-2 nucleocapsid protein regulates RNA binding, its ability to suppress stress granule formation, and viral replication. Journal of Biological Chemistry. 2021;297: 100821. doi:10.1016/j.jbc.2021.100821

52. Luo L, Li Z, Zhao T, Ju X, Ma P, Jin B, et al. SARS-CoV-2 nucleocapsid protein phase separates with G3BPs to disassemble stress granules and facilitate viral production. Sci Bull (Beijing). 2021;66: 1194–1204. doi:10.1016/j.scib.2021.01.013

53. Mulloy RP, Evseev D, Sharlin N, Bui-Marinos MP, Lacasse É, Dubuc I, et al. Evolution of a truncated nucleocapsid protein enhances SARS-CoV-2 fitness by suppressing antiviral responses. PLOS Biology. 2026;24: e3003646. doi:10.1371/journal.pbio.3003646

54. Nabeel-Shah S, Lee H, Ahmed N, Burke GL, Farhangmehr S, Ashraf K, et al. SARS-CoV-2 nucleocapsid protein binds host mRNAs and attenuates stress granules to impair host stress response. iScience. 2022;25: 103562. doi:10.1016/j.isci.2021.103562

55. Yang Z, Johnson BA, Meliopoulos VA, Ju X, Zhang P, Hughes MP, et al. Interaction between host G3BP and viral nucleocapsid protein regulates SARS-CoV-2 replication and pathogenicity. Cell Reports. 2024;43. doi:10.1016/j.celrep.2024.113965

56. Zheng Y, Deng J, Han L, Zhuang M-W, Xu Y, Zhang J, et al. SARS-CoV-2 NSP5 and N protein counteract the RIG-I signaling pathway by suppressing the formation of stress granules. Sig Transduct Target Ther. 2022;7: 22. doi:10.1038/s41392-022-00878-3

57. Zheng Z-Q, Wang S-Y, Xu Z-S, Fu Y-Z, Wang Y-Y. SARS-CoV-2 nucleocapsid protein impairs stress granule formation to promote viral replication. Cell Discov. 2021;7: 1–11. doi:10.1038/s41421-021-00275-0

58. Cui L, Wang H, Ji Y, Yang J, Xu S, Huang X, et al. The Nucleocapsid Protein of Coronaviruses Acts as a Viral Suppressor of RNA Silencing in Mammalian Cells. Journal of Virology. 2015;89: 9029–9043. doi:10.1128/jvi.01331-15

59. LeBlanc K, Lynch J, Layne C, Vendramelli R, Sloan A, Tailor N, et al. The Nucleocapsid Proteins of SARS-CoV-2 and Its Close Relative Bat Coronavirus RaTG13 Are Capable of Inhibiting PKR- and RNase L-Mediated Antiviral Pathways. Microbiol Spectr. 2023;11: e0099423. doi:10.1128/spectrum.00994-23

60. Lodola C, Secchi M, Sinigiani V, De Palma A, Rossi R, Perico D, et al. Interaction of SARS-CoV-2 Nucleocapsid Protein and Human RNA Helicases DDX1 and DDX3X Modulates Their Activities on Double-Stranded RNA. International Journal of Molecular Sciences. 2023;24: 5784. doi:10.3390/ijms24065784

61. Adly AN, Bi M, Carlson CR, Syed AM, Ciling A, Doudna JA, et al. Assembly of SARS-CoV-2 ribonucleosomes by truncated N∗ variant of the nucleocapsid protein. Journal of Biological Chemistry. 2023;299. doi:10.1016/j.jbc.2023.105362

62. Mears HV, Young GR, Sanderson T, Harvey R, Barrett-Rodger J, Penn R, et al. Emergence of SARS-CoV-2 subgenomic RNAs that enhance viral fitness and immune evasion. PLOS Biology. 2025;23: e3002982. doi:10.1371/journal.pbio.3002982

63. Syed AM, Ciling A, Chen IP, Carlson CR, Adly AN, Martin HS, et al. SARS-CoV-2 evolution balances conflicting roles of N protein phosphorylation. PLOS Pathogens. 2024;20: e1012741. doi:10.1371/journal.ppat.1012741

64. Edgell CJ, McDonald CC, Graham JB. Permanent cell line expressing human factor VIII-related antigen established by hybridization. Proceedings of the National Academy of Sciences. 1983;80: 3734–3737. doi:10.1073/pnas.80.12.3734

65. Dolliver SM, Galbraith C, Khaperskyy DA. Human Betacoronavirus OC43 Interferes with the Integrated Stress Response Pathway in Infected Cells. Viruses. 2024;16: 212. doi:10.3390/v16020212

66. Renner DM, Parenti NA, Bracci N, Weiss SR. Betacoronaviruses Differentially Activate the Integrated Stress Response to Optimize Viral Replication in Lung-Derived Cell Lines. Viruses. 2025;17: 120. doi:10.3390/v17010120

67. Shaban MS, Weiser HS, Weber A, Meier-Soelch J, Dort F, Mayr-Buro C, et al. PERK inhibition rewires translational and CMGC protein kinase networks into an antiviral state. bioRxiv; 2025. p. 2025.10.08.681090. doi:10.1101/2025.10.08.681090

68. Shaban MS, Müller C, Mayr-Buro C, Weiser H, Meier-Soelch J, Albert BV, et al. Multi-level inhibition of coronavirus replication by chemical ER stress. Nat Commun. 2021;12: 5536. doi:10.1038/s41467-021-25551-1

69. Davies EL, Sowar H, Balci A, Moorhouse E, Wickenhagen A, Turnbull ML, et al. Alternative splicing broadens antiviral diversity at the human OAS2 locus. bioRxiv; 2025. p. 2025.02.24.639105. doi:10.1101/2025.02.24.639105

70. Merold V, Bekere I, Kretschmer S, Schnell AF, Kmiec D, Sivarajan R, et al. Structural basis for OAS2 regulation and its antiviral function. Molecular Cell. 2025;85: 2176–2193.e13. doi:10.1016/j.molcel.2025.05.001

71. Soveg FW, Schwerk J, Gokhale NS, Cerosaletti K, Smith JR, Pairo-Castineira E, et al. Endomembrane targeting of human OAS1 p46 augments antiviral activity. Van der Meer JW, editor. eLife. 2021;10: e71047. doi:10.7554/eLife.71047

72. Wickenhagen A, Sugrue E, Lytras S, Kuchi S, Noerenberg M, Turnbull ML, et al. A prenylated dsRNA sensor protects against severe COVID-19. Science. 2021;374: eabj3624. doi:10.1126/science.abj3624

73. Wang J, Shi C, Xu Q, Yin H. SARS-CoV-2 nucleocapsid protein undergoes liquid–liquid phase separation into stress granules through its N-terminal intrinsically disordered region. Cell Discov. 2021;7: 1–5. doi:10.1038/s41421-020-00240-3

74. Peng T-Y, Lee K-R, Tarn W-Y. Phosphorylation of the arginine/serine dipeptide-rich motif of the severe acute respiratory syndrome coronavirus nucleocapsid protein modulates its multimerization, translation inhibitory activity and cellular localization. The FEBS Journal. 2008;275: 4152–4163. doi:10.1111/j.1742-4658.2008.06564.x

75. Long S, Guzyk M, Perez Vidakovics L, Han X, Sun R, Wang M, et al. SARS-CoV-2 N protein recruits G3BP to double membrane vesicles to promote translation of viral mRNAs. Nat Commun. 2024;15: 10607. doi:10.1038/s41467-024-54996-3

76. Alvarado RE, Chen J, Lokugamage KG, Zhou Y, Estes LK, Morgan A, et al. Antagonism of stress granules key for SARS-CoV-2 infection and pathogenesis. bioRxiv; 2026. p. 2026.06.16.732644. doi:10.64898/2026.06.16.732644

77. Deng X, Hackbart M, Mettelman RC, O’Brien A, Mielech AM, Yi G, et al. Coronavirus nonstructural protein 15 mediates evasion of dsRNA sensors and limits apoptosis in macrophages. Proceedings of the National Academy of Sciences. 2017;114: E4251–E4260. doi:10.1073/pnas.1618310114

78. Gong X, Feng S, Wang J, Gao B, Xue W, Chu H, et al. Coronavirus endoribonuclease nsp15 suppresses host protein synthesis and evades PKR-eIF2α-mediated translation shutoff to ensure viral protein synthesis. PLOS Pathogens. 2025;21: e1012987. doi:10.1371/journal.ppat.1012987

79. Kindler E, Gil-Cruz C, Spanier J, Li Y, Wilhelm J, Rabouw HH, et al. Early endonuclease-mediated evasion of RNA sensing ensures efficient coronavirus replication. PLOS Pathogens. 2017;13: e1006195. doi:10.1371/journal.ppat.1006195

80. Goldstein SA, Thornbrough JM, Zhang R, Jha BK, Li Y, Elliott R, et al. Lineage A Betacoronavirus NS2 Proteins and the Homologous Torovirus Berne pp1a Carboxy-Terminal Domain Are Phosphodiesterases That Antagonize Activation of RNase L. J Virol. 2017;91: e02201–16. doi:10.1128/JVI.02201-16

81. Thornbrough JM, Jha BK, Yount B, Goldstein SA, Li Y, Elliott R, et al. Middle East Respiratory Syndrome Coronavirus NS4b Protein Inhibits Host RNase L Activation. mBio. 2016;7: 10.1128/mbio.00258-16. doi:10.1128/mbio.00258-16

82. Bhardwaj K, Guarino L, Kao CC. The Severe Acute Respiratory Syndrome Coronavirus Nsp15 Protein Is an Endoribonuclease That Prefers Manganese as a Cofactor. J Virol. 2004;78: 12218–12224. doi:10.1128/JVI.78.22.12218-12224.2004

83. Hackbart M, Deng X, Baker SC. Coronavirus endoribonuclease targets viral polyuridine sequences to evade activating host sensors. Proceedings of the National Academy of Sciences. 2020;117: 8094–8103. doi:10.1073/pnas.1921485117

84. Wang X, Zhu B. SARS-CoV-2 nsp15 preferentially degrades AU-rich dsRNA via its dsRNA nickase activity. Nucleic Acids Res. 2024;52: 5257–5272. doi:10.1093/nar/gkae290

85. Otter CJ, Renner DM, Fausto A, Tan LH, Cohen NA, Weiss SR. Interferon signaling in the nasal epithelium distinguishes among lethal and common cold coronaviruses and mediates viral clearance. Proceedings of the National Academy of Sciences. 2024;121: e2402540121. doi:10.1073/pnas.2402540121

86. Li Y, Renner DM, Comar CE, Whelan JN, Reyes HM, Cardenas-Diaz FL, et al. SARS-CoV-2 induces double-stranded RNA-mediated innate immune responses in respiratory epithelial-derived cells and cardiomyocytes. Proceedings of the National Academy of Sciences. 2021;118: e2022643118. doi:10.1073/pnas.2022643118

87. Dong H, Zhang H, Jalin J, He Z, Wang R, Huang L, et al. Nucleocapsid proteins from human coronaviruses possess phase separation capabilities and promote FUS pathological aggregation. Protein Science. 2023;32: e4826. doi:10.1002/pro.4826

88. Ahmad S, Zou T, Hwang J, Zhao L, Wang X, Davydenko A, et al. PACT prevents aberrant activation of PKR by endogenous dsRNA without sequestration. Nat Commun. 2025;16: 3325. doi:10.1038/s41467-025-58433-x

89. Corbet GA, Burke JM, Parker R. Nucleic acid–protein condensates in innate immune signaling. The EMBO Journal. 2023;42: e111870. doi:10.15252/embj.2022111870

90. He S, Gou H, Zhou Y, Wu C, Ren X, Wu X, et al. The SARS-CoV-2 nucleocapsid protein suppresses innate immunity by remodeling stress granules to atypical foci. The FASEB Journal. 2023;37: e23269. doi:10.1096/fj.202201973RR

91. Carlson CR, Adly AN, Bi M, Howard CJ, Frost A, Cheng Y, et al. Reconstitution of the SARS-CoV-2 ribonucleosome provides insights into genomic RNA packaging and regulation by phosphorylation. Journal of Biological Chemistry. 2022;298. doi:10.1016/j.jbc.2022.102560

92. Carlson CR, Asfaha JB, Ghent CM, Howard CJ, Hartooni N, Safari M, et al. Phosphoregulation of Phase Separation by the SARS-CoV-2 N Protein Suggests a Biophysical Basis for its Dual Functions. Mol Cell. 2020;80: 1092–1103.e4. doi:10.1016/j.molcel.2020.11.025

93. Lu S, Ye Q, Singh D, Cao Y, Diedrich JK, Yates JR, et al. The SARS-CoV-2 nucleocapsid phosphoprotein forms mutually exclusive condensates with RNA and the membrane-associated M protein. Nat Commun. 2021;12: 502. doi:10.1038/s41467-020-20768-y

94. Ren J, Wang S, Zong Z, Pan T, Liu S, Mao W, et al. TRIM28-mediated nucleocapsid protein SUMOylation enhances SARS-CoV-2 virulence. Nat Commun. 2024;15: 244. doi:10.1038/s41467-023-44502-6

95. Rhamadianti AF, Abe T, Tanaka T, Ono C, Katayama H, Makino Y, et al. SARS-CoV-2 papain-like protease inhibits ISGylation of the viral nucleocapsid protein to evade host anti-viral immunity. Journal of Virology. 2024;98: e00855–24. doi:10.1128/jvi.00855-24

96. Stuwe H, Reardon PN, Yu Z, Shah S, Hughes K, Barbar EJ. Phosphorylation in the Ser/Arg-rich region of the nucleocapsid of SARS-CoV-2 regulates phase separation by inhibiting self-association of a distant helix. Journal of Biological Chemistry. 2024;300. doi:10.1016/j.jbc.2024.107354

97. Wu C-H, Chen P-J, Yeh S-H. Nucleocapsid Phosphorylation and RNA Helicase DDX1 Recruitment Enables Coronavirus Transition from Discontinuous to Continuous Transcription. Cell Host & Microbe. 2014;16: 462–472. doi:10.1016/j.chom.2014.09.009

98. Yaron TM, Heaton BE, Levy TM, Johnson JL, Jordan TX, Cohen BM, et al. Host protein kinases required for SARS-CoV-2 nucleocapsid phosphorylation and viral replication. Sci Signal. 2022;15: eabm0808. doi:10.1126/scisignal.abm0808

99. Zhu J, Liu G, Sayyad Z, Goins CM, Stauffer SR, Gack MU. ISGylation of the SARS-CoV-2 N protein by HERC5 impedes N oligomerization and thereby viral RNA synthesis. Journal of Virology. 2024;98: e00869–24. doi:10.1128/jvi.00869-24

100. Tanneti NS, Patel AK, Tan LH, Marques AD, Perera RAPM, Sherrill-Mix S, et al. Comparison of SARS-CoV-2 variants of concern in primary human nasal cultures demonstrates Delta as most cytopathic and Omicron as fastest replicating. mBio. 15: e03129–23. doi:10.1128/mbio.03129-23

101. Gartner MJ, Smith ML, Dapat C, Liaw YW, Tran T, Suryadinata R, et al. Contemporary seasonal human coronaviruses display differences in cellular tropism compared to laboratory-adapted reference strains. Journal of Virology. 2025;99: e00684–25. doi:10.1128/jvi.00684-25

102. Ye Y, Hauns K, Langland JO, Jacobs BL, Hogue BG. Mouse Hepatitis Coronavirus A59 Nucleocapsid Protein Is a Type I Interferon Antagonist. Journal of Virology. 2007;81: 2554–2563. doi:10.1128/JVI.01634-06

103. Borgogna C, Cislaghi I, Turati S, Mozzi A, Forni D, Cagliani R, et al. Convergent evolution of the G3BP1-binding motif in betacoronavirus nucleocapsid proteins. Virus Evol. 2025;11: veaf063. doi:10.1093/ve/veaf063

104. Kleer M, Mulloy RP, Robinson C-A, Evseev D, Bui-Marinos MP, Castle EL, et al. Human coronaviruses disassemble processing bodies. PLOS Pathogens. 2022;18: e1010724. doi:10.1371/journal.ppat.1010724

105. Gordon DE, Jang GM, Bouhaddou M, Xu J, Obernier K, White KM, et al. A SARS-CoV-2 protein interaction map reveals targets for drug repurposing. Nature. 2020;583: 459–468. doi:10.1038/s41586-020-2286-9

106. Robinson C-A, Singh GK, Kleer M, Katsademas T, Castle EL, Boudreau BQ, et al. Kaposi’s sarcoma-associated herpesvirus (KSHV) utilizes the NDP52/CALCOCO2 selective autophagy receptor to disassemble processing bodies. PLOS Pathogens. 2023;19: e1011080. doi:10.1371/journal.ppat.1011080

107. Stirling DR, Swain-Bowden MJ, Lucas AM, Carpenter AE, Cimini BA, Goodman A. CellProfiler 4: improvements in speed, utility and usability. BMC Bioinformatics. 2021;22: 433. doi:10.1186/s12859-021-04344-9

108. Madeira F, Madhusoodanan N, Lee J, Eusebi A, Niewielska A, Tivey ARN, et al. The EMBL-EBI Job Dispatcher sequence analysis tools framework in 2024. Nucleic Acids Res. 2024;52: W521–W525. doi:10.1093/nar/gkae241

109. Sievers F, Wilm A, Dineen D, Gibson TJ, Karplus K, Li W, et al. Fast, scalable generation of high-quality protein multiple sequence alignments using Clustal Omega. Mol Syst Biol. 2011;7: 539. doi:10.1038/msb.2011.75

110. Waterhouse AM, Procter JB, Martin DMA, Clamp M, Barton GJ. Jalview Version 2—a multiple sequence alignment editor and analysis workbench. Bioinformatics. 2009;25: 1189–1191. doi:10.1093/bioinformatics/btp033

111. Blum M, Andreeva A, Florentino LC, Chuguransky SR, Grego T, Hobbs E, et al. InterPro: the protein sequence classification resource in 2025. Nucleic Acids Res. 2025;53: D444–D456. doi:10.1093/nar/gkae1082

