## Supplemental Figs for "Human coronavirus nucleocapsid proteins have disparate innate immune evasion abilities"

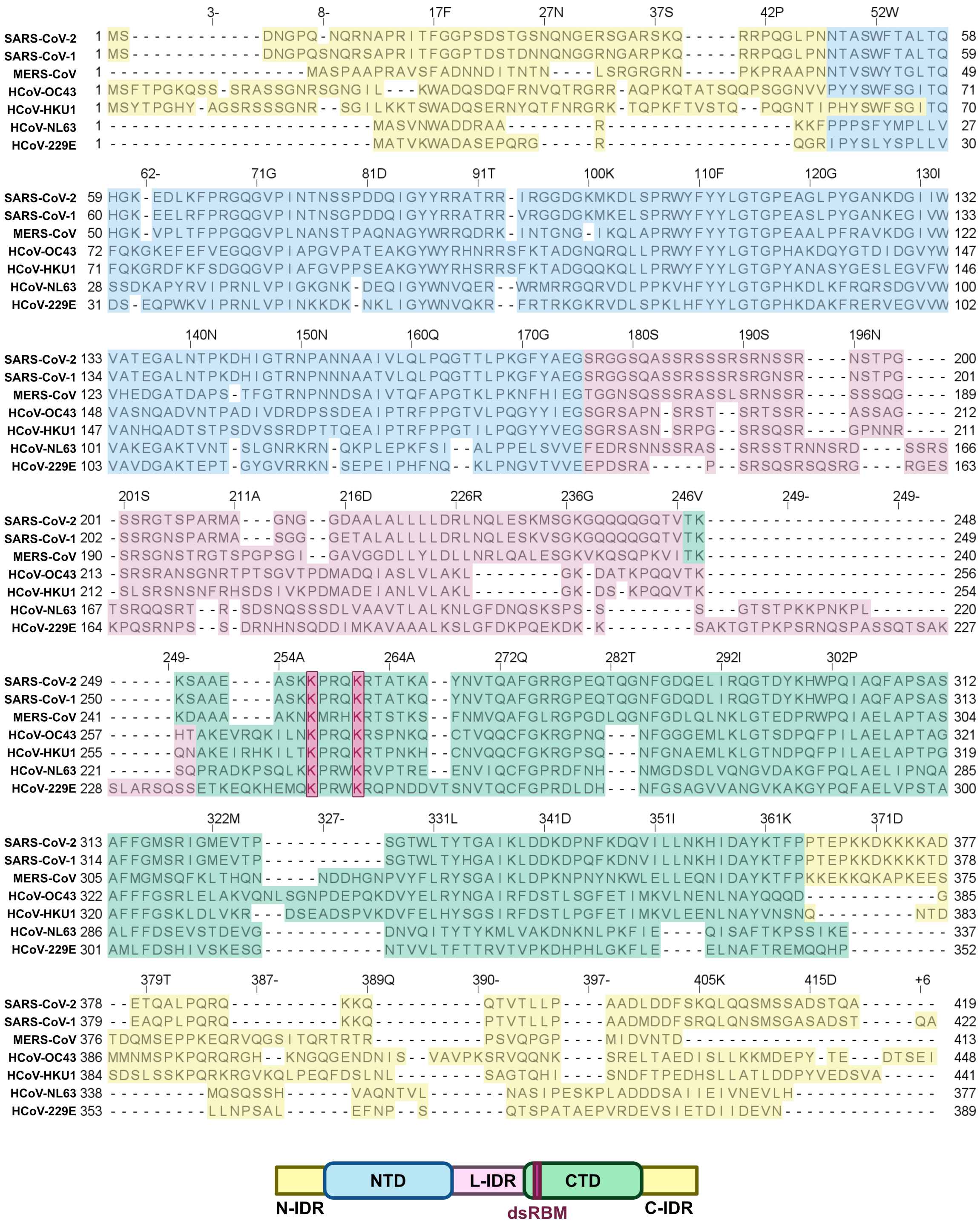
SARS-CoV-2
SARS-CoV-1
MERS-CoV
HCoV-OC43
HCoV-HKU1
HCoV-NL63
HCoV-229E
NTD
L-IDR
CTD
N-IDR
dsRBM
C-IDR

**Figure S1. Multiple sequence alignment of HCoV N proteins.**

HCoV N protein sequences were aligned using Clustal Omega, and domain boundaries were delineated using InterPro predictions. Residue numbering for SARS-CoV-2 N is depicted above the top row of the alignment. Residues are colour-coded by domain as shown by the domain diagram below. dsRNA-binding motif (dsRBM) lysines are indicated with purple boxes.

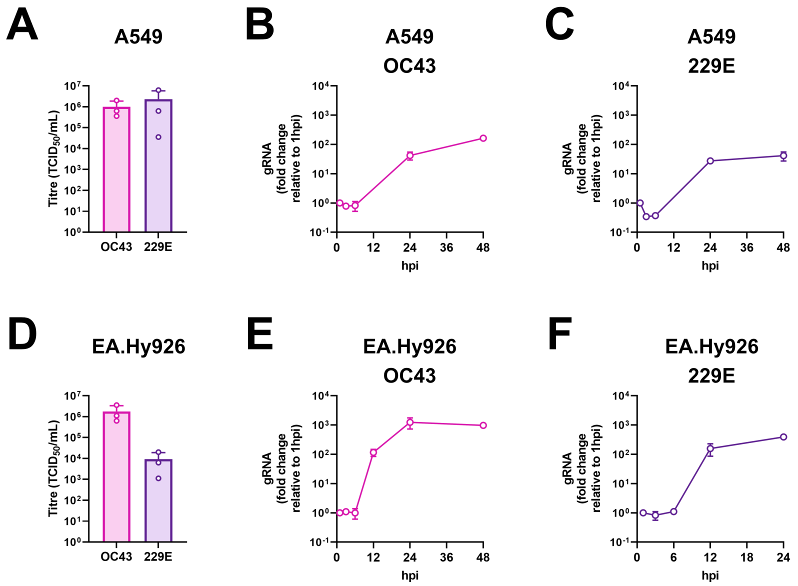
A
A549
Titre (TCID50/mL)
OC43
229E
B
A549
OC43
gRNA (fold change relative to 1hpi)
hpi
C
A549
229E
gRNA (fold change relative to 1hpi)
hpi
D
EA.Hy926
Titre (TCID50/mL)
OC43
229E
E
EA.Hy926
OC43
gRNA (fold change relative to 1hpi)
hpi
F
EA.Hy926
229E
gRNA (fold change relative to 1hpi)
hpi

**Figure S2. EA.Hy926 cells are permissive to HCoV-OC43 and HCoV-229E.**

**A.** A549 cells were infected with HCoV-OC43 or HCoV-229E at an MOI of 1. Virus-containing supernatants were harvested at 48 hpi for HCoV-OC43 or 24 hpi for HCoV-229E, and viral titres were determined by $TCID_{50}$. These data represent three independent biological replicates ($n$ = 3; mean + SD).

**B-C.** A549 cells were infected with HCoV-OC43 (B) or HCoV-229E (C) at an MOI of 1. Intracellular RNA was harvested at indicated times post-infection and viral gRNA abundance was measured by RT-qPCR. Data are represented as fold change relative to 1 hpi from three independent biological replicates ($n$ = 3; mean ± SD).

**D.** EA.Hy926 cells were infected with HCoV-OC43 or HCoV-229E at an MOI of 1 and viral titres were determined as in A. These data represent three independent biological replicates ($n$ = 3; mean ± SD).

**E-F.** EA.Hy926 cells were infected with HCoV-OC43 (E) or HCoV-229E (F) at an MOI of 1 and viral gRNA abundance was measured as in B-C. Data are represented as fold change relative to 1 hpi from three independent biological replicates ($n$ = 3; mean ± SD).

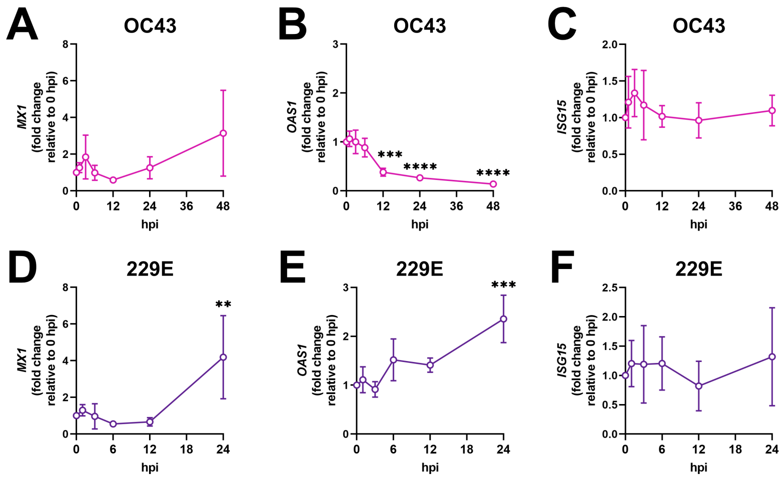
A
OC43
MX1
(fold change
relative to 0 hpi)
hpi
B
OC43
OAS1
(fold change
relative to 0 hpi)
***
****
****
hpi
C
OC43
ISG15
(fold change
relative to 0 hpi)
hpi
D
229E
MX1
(fold change
relative to 0 hpi)
**
hpi
E
229E
OAS1
(fold change
relative to 0 hpi)
***
hpi
F
229E
ISG15
(fold change
relative to 0 hpi)
hpi

**Figure S3. HCoV-OC43 and HCoV-229E induce distinct ISG expression profiles during infection.**

**A-F.** EA.Hy926 cells were infected with HCoV-OC43 (A-C) or HCoV-229E (D-F) at an MOI of 1. Intracellular RNA was harvested at indicated times post-infection and subjected to RT-qPCR to measure the abundance of *MX1* (A, D), *OAS1* (B, E), and *ISG15* (C, F) mRNAs. Data are represented as fold change relative to 0 hpi from three independent biological replicates ($n$ = 3; mean ± SD). Statistics were performed relative to 0 hpi using a one-way ANOVA with Dunnett's post-hoc analysis (****, $p < 0.0001$; ***, $p < 0.001$; **, $p < 0.01$; no symbol shown, nonsignificant).

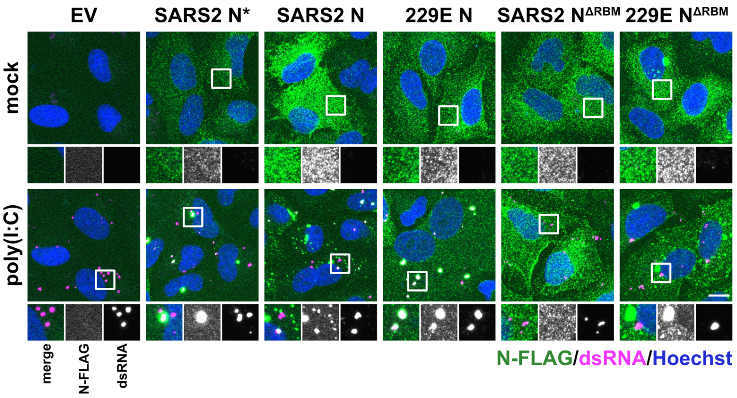

EV
SARS2 N*
SARS2 N
229E N
SARS2 N^ΔRBM
229E N^ΔRBM
mock
poly(I:C)
merge
N-FLAG
dsRNA
N-FLAG/dsRNA/Hoechst

**Figure S4. SARS-CoV-2 and HCoV-229E N proteins colocalize with dsRNA in a dsRNA-binding motif (dsRBM)-dependent manner.**

A549 cells were transduced with recombinant lentiviruses expressing an empty vector control (EV) or each of the indicated HCoV N-FLAG constructs and selected with puromycin. The dsRBM lysines were mutated to alanine for the $SARS2^{\Delta RBM}$ construct (K257A/K261A) or arginine for the $229E^{\Delta RBM}$ construct (K246R/K250R). Six days post-transduction, cells were mock-transfected or transfected with 0.5 µg poly(I:C). Cells were fixed three hours post-poly(I:C) addition and immunostained with antibodies specific to FLAG (N protein; green) or dsRNA (magenta). Nuclei were stained with Hoechst (blue). Images were obtained using an LSM 880 Airyscan confocal microscope, with maximum-intensity projections presented (scale bar = 10 µm). The images shown represent one of two independent biological replicates ($n = 2$).

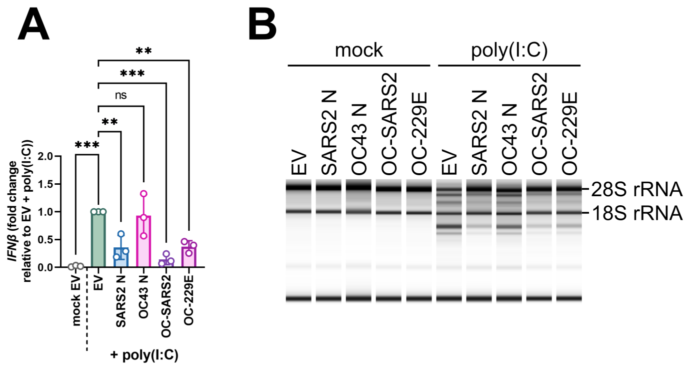
A
IFNβ (fold change relative to EV + poly(I:C))
mock EV
EV
SARS2 N
OC43 N
OC-SARS2
OC-229E
+ poly(I:C)
B
mock
poly(I:C)
EV
SARS2 N
OC43 N
OC-SARS2
OC-229E
28S rRNA
18S rRNA
C
rRNA integrity (RIN^e)
mock EV
EV
SARS2 N
OC43 N
OC-SARS2
OC-229E
+ poly(I:C)

**Figure S5. The carboxy termini of HCoV N proteins modulate their interferon and OAS/RNase L evasion abilities.**

**A.** A549 cells were transduced with recombinant lentiviruses expressing an empty vector control (EV) or each of the indicated HCoV N-FLAG constructs and selected with puromycin. Six days post-transduction, cells were mock-transfected or transfected with 0.5 μg poly(I:C). Intracellular RNA was harvested three hours post-poly(I:C) addition and subjected to RT-qPCR to measure *IFNβ* mRNA abundance. Values were normalized to 18S rRNA (housekeeping control) and are represented as fold change relative to the EV-transduced, poly(I:C)-transfected condition from three independent biological replicates ($n$ = 3; mean ± SD). Statistics were performed using a one-way ANOVA with Dunnett's post-hoc analysis (***, $p < 0.001$; **, $p < 0.01$; ns, nonsignificant).

**B.** A549 cells were transduced, selected, and transfected with poly(I:C) as in A. Intracellular RNA was harvested three hours post-poly(I:C) addition and subjected to automated electrophoresis using an Agilent 4200 TapeStation system to assess rRNA integrity. The electronic gel shown represents one of three independent biological replicates ($n$ = 3).

**C.** The RNA integrity number equivalent ($RIN^e$) was measured for the samples analyzed by automated electrophoresis in B. These data represent three independent biological replicates ($n$ = 3; mean ± SD). Statistics were performed using a one-way ANOVA with Dunnett's post-hoc analysis (***, $p < 0.001$; **, $p < 0.01$; ns, nonsignificant).
